# Systemic ablation of *Camkk2* impairs metastatic colonization and improves insulin sensitivity in TRAMP mice: Evidence for cancer cell-extrinsic CAMKK2 functions in prostate cancer

**DOI:** 10.1101/2022.03.28.486100

**Authors:** Thomas L. Pulliam, Dominik Awad, Jenny J. Han, Mollianne M. Murray, Jeffrey J. Ackroyd, Pavithr Goli, Jonathan S. Oakhill, John W. Scott, Michael M. Ittman, Daniel E. Frigo

## Abstract

Despite early studies linking calcium-calmodulin protein kinase kinase 2 (CAMKK2) to prostate cancer cell migration and invasion, the role of CAMKK2 in metastasis *in vivo* remains unclear. Moreover, while CAMKK2 is known to regulate systemic metabolism, whether CAMKK2’s effects on whole body metabolism would impact prostate cancer progression and/or related comorbidities is not known. Here, we demonstrate that germline ablation of *Camkk2* slows, but does not stop, primary prostate tumorigenesis in the <u>TR</u>ansgenic <u>A</u>denocarcinoma <u>M</u>ouse <u>P</u>rostate (TRAMP) genetic mouse model. Consistent with prior epidemiological reports supporting a link between obesity and prostate cancer aggressiveness, TRAMP mice fed a high-fat diet exhibited a pronounced increase in the colonization of lung metastases. We demonstrated that this effect on metastatic spread was dependent on CAMKK2. Notably, diet-induced lung metastases exhibited a highly aggressive neuroendocrine phenotype. Concurrently, *Camkk2* deletion improved insulin sensitivity in the same mice. Histological analyses revealed that cancer cells were smaller in the TRAMP;*Camkk2*-/- mice compared to TRAMP;*Camkk2*+/+ controls. Given the differences in circulating insulin levels, a known regulator of cell growth, we hypothesized that systemic CAMKK2 could promote prostate cancer cell growth and disease progression in part through cancer cell-extrinsic mechanisms. Accordingly, host deletion of *Camkk2* impaired the growth of syngeneic murine prostate tumors *in vivo*, confirming nonautonomous roles for CAMKK2 in prostate cancer. Cancer cell size and mTOR signaling was diminished in tumors propagated in *Camkk2*-null mice. Together, these data indicate that, in addition to cancer cell-intrinsic roles, CAMKK2 promotes prostate cancer progression via tumor-extrinsic mechanisms. Further, we propose that CAMKK2 inhibition may also help combat common metabolic comorbidities in men with advanced prostate cancer.

## Introduction

Prostate cancer is the most commonly diagnosed non-skin cancer in men worldwide [1]. If detected early, prostate cancer is treatable with surgery and/or radiation. However, if allowed to proceed, the disease becomes incurable. Because of its high prevalence, prostate cancer accounts for the second most cancer-related deaths in the U.S. [1]

Patients presenting with prostate cancer that cannot be surgically resected or treated with local radiation are initially given androgen deprivation therapy (ADT). ADT inhibits activation of the transcription factor and primary driver of prostate cancer, the androgen receptor (AR) [2]. While initially effective, the disease often relapses within 2-3 years, resulting in a form of the disease called castration-resistant prostate cancer (CRPC) that ironically remains largely driven by AR. Subsequent second-generation AR antagonists and other non-AR targeting therapies have been approved for the treatment of CRPC, but these treatments are not curative. Due to the resistance associated with ADT and second-generation AR-targeting drugs, it is vital to identify alternative therapeutic targets driving CRPC downstream of reactivated AR. One direct AR downstream target that promotes prostate cancer progression is the calcium-calmodulin-dependent protein kinase kinase 2 (CAMKK2) [3-5].

CAMKK2 is a serine/threonine kinase that facilitates cancer progression via the phosphorylation and activation of downstream targets such as CAMKI, CAMKIV, and AMPK [6-8]. Correspondingly, treatment with the CAMKK2 inhibitor STO-609 decreased tumor growth in a CRPC subcutaneous xenograft mouse model, demonstrating that CAMKK2 can be pharmacologically targeted [4]. Previous work using a germline knockout (KO) of *Camkk2* in the Pb-Cre;*Pten*^f/f^ genetic mouse model (GEMM) of prostate cancer demonstrated decreased incidence and severity of prostatic intraepithelial neoplasia (PIN) compared to *Camkk2* WT mice [9]. Importantly, *Camkk2* ablation blocked progression to primary adenocarcinoma. However, the Pb-Cre;*Pten*^f/f^ GEMM, as a relatively benign prostate cancer model, does not allow CAMKK2 to be studied in the context of metastasis or progression to highly aggressive subtypes such as neuroendocrine prostate cancer (NEPC). Because CAMKK2 is associated with prostate cancer migration and invasion *in vitro* [3], we utilized the more aggressive GEMM, <u>TR</u>ansgenic <u>A</u>denocarcinoma <u>M</u>ouse <u>P</u>rostate (TRAMP), which can recapitulate multiple stages of prostate cancer progression including metastasis and transition to NEPC [10,11], to explore CAMKK2’s role in advanced prostate cancer *in vivo*.

Beyond CAMKK2’s role in cancer, CAMKK2 regulates several other physiological conditions that could indirectly contribute to cancer progression. Inhibition of CAMKK2 enhances immune response to infection, promotes macrophage maturation, and regresses age-related bone degeneration [12-16]. In addition, CAMKK2 inhibition protects mice from developing metabolic syndrome shown through a decrease in blood glucose and insulin resistance as well as regression of nonalcoholic fatty liver disease (NAFLD) [17-19]. This phenomenon was of interest to us because a major comorbidity for prostate cancer patients is an increased risk of metabolic syndrome [20,21]. Thus, in addition to testing CAMKK2’s role in an aggressive prostate cancer GEMM, we sought to simultaneously assess its impact on systemic metabolism to determine if targeting CAMKK2 could be dually beneficial in both treating the tumor and its linked comorbidities.

## Materials and Methods

### Animal experiments

All animal experiments were approved by and conducted under the Institutional Animal Care and Use Committee at the University of Texas MD Anderson Cancer Center (MDACC) and the University of Houston (UH) according to NIH and institutional guidelines (MDACC IACUC #00001738-RN01 and UH IACUC #14-039).

### GEMM models

Original *Camkk2*-/- mice have previously been described [19]. C57BL/6 TRAMP mice (Strain #003135) were obtained from Jackson Laboratories. To create a germline *Camkk2* knockout line for syngeneic experiments and one that could be more directly comparable to existing *Camkk2* floxed mice for future studies (note, *Camkk2*-/- mice crossed to TRAMP mice in this study were original *Camkk2*-/- mice while *Camkk2*-/- mice used in syngeneic experiments were created from *Camkk2* floxed mice [22], we crossed female Sox2-Cre mice (mixed background) to male *Camkk2*^f/f^ mice (C57BL/6J) to create *Camkk2-*null allele-carrying mice. This approach caused the expression of Cre in early oogenesis prior to any differentiation, allowing floxed alleles to be deleted after the fusion of the pronuclei in the fertilized egg in what would become embryonic germ cells [23]. Germline knockouts/null alleles were then detected with genotyping and selected for further breeding. Founder mice were backcrossed onto a pure C57BL6/J genetic background for >7 generations. Single nucleotide polymorphism (SNP) analysis was carried out by the MDACC Laboratory Animal Genetic Services Core to characterize the background strain purity. Newly derived *Camkk2*-/- mice were confirmed to be 99-99.5% C57BL/6, whereas the original *Camkk2*-/- mice were 80.5-85% C57BL/6. Immunoblot analysis further confirmed the loss of CAMKK2 protein in the newly derived mice (Supplemental Figure S1).

### Immunoblot analysis

Pieces of mouse brains (specifically cerebellum) were resected and homogenized using a handheld homogenizer in 1 mL ice-cold RIPA buffer and agitated at 4°C for 1 hour. Protein concentrations were measured using Bradford’s reagent. Immunoblotting was conducted as before [24].

### High-fat diet studies

High-fat diet (HFD) was obtained from Bio-serv (60% kcals from fat, cat#: S3282). TRAMP*;Camkk2+/+* and TRAMP*;Camkk2-/-* mice (described in *GEMM Models*) were fed control chow until 6 weeks of age and then stayed on control chow or switched to HFD for 15 or 30 weeks. At endpoints, prostates/tumors, kidneys, and livers were removed and weighed for all mice.

### Plasma

Plasma was collected in heparin tubes by cardiac puncture and analyzed for circulating glucose, insulin, triglycerides, cholesterol, free fatty acids, and leptin by the Baylor College of Medicine (BCM) Mouse Metabolism and Phenotyping Core. <u>HO</u>meostatic <u>M</u>odel <u>A</u>ssessment for Insulin <u>R</u>esistance* (HOMA-IR*) was calculated using the formula: (glucose x insulin)/22.5. In this study, mice were not fasted prior to the collection of the plasma.

### Cell culture

RM-9 cells were gifted from the laboratory of Timothy Thompson at MDACC and grown in Dulbecco’s Modified Eagle Medium (DMEM) supplemented with 10% FBS, 200 mM L-glutamine, and 1 M HEPES. TRAMP-C2 cells were attained from ATCC and grown in DMEM supplemented with 5% FBS, 5% Nu-Serum (VWR Cat#45001-090), insulin (cell culture grade) (VWR Cat# 47743-634), and 0.01 nM dihydrotestosterone (DHT).

### Proliferation assays

Proliferation assays were carried out as previously described [25]. Briefly, 5,000 RM-9 or TRAMP-C2 cells were plated in charcoal-stripped serum for 72 hours and then treated with vehicle or indicated androgens for 7 days and relative cell numbers were quantified using a Hochest-based DNA dye.

### Syngeneic models

50,000 RM-9 cells in 200 µl DPBS were injected subcutaneously into flanks of 8-week-old, *Camkk2+/+* and *Camkk2-/-* mice. 1×10^6^ TRAMP-C2 cells in 200 µl DMEM: Matrigel® (Corning, Corning, NY, USA; Cat #356231) 1:1 vol/vol were injected subcutaneously into flanks of 8-week-old, *Camkk2*+/+ and *Camkk2-/-* mice. Tumor volume was calculated by the formula: (length x width^2^)/2. Mice were harvested when tumors reached 1.5 cm^3^.

### Histology and immunostaining

Processing, embedding, hematoxylin and eosin (H&E), and immunostaining were conducted by the MD Anderson Cancer Center DVMS Veterinary Pathology Services (Houston, TX, USA). Additional processing, embedding, H&E, and immunostaining were conducted by the MD Anderson Cancer Center Science Park Research Histology, Pathology and Imaging Core (RHPI) (Smithville, TX, USA). Additional immunostaining was conducted by HistoWiz (Brooklyn, NY, USA). Image analysis was conducted using QuPath v0.3.0.

### Antibody Table

**Table 1.**
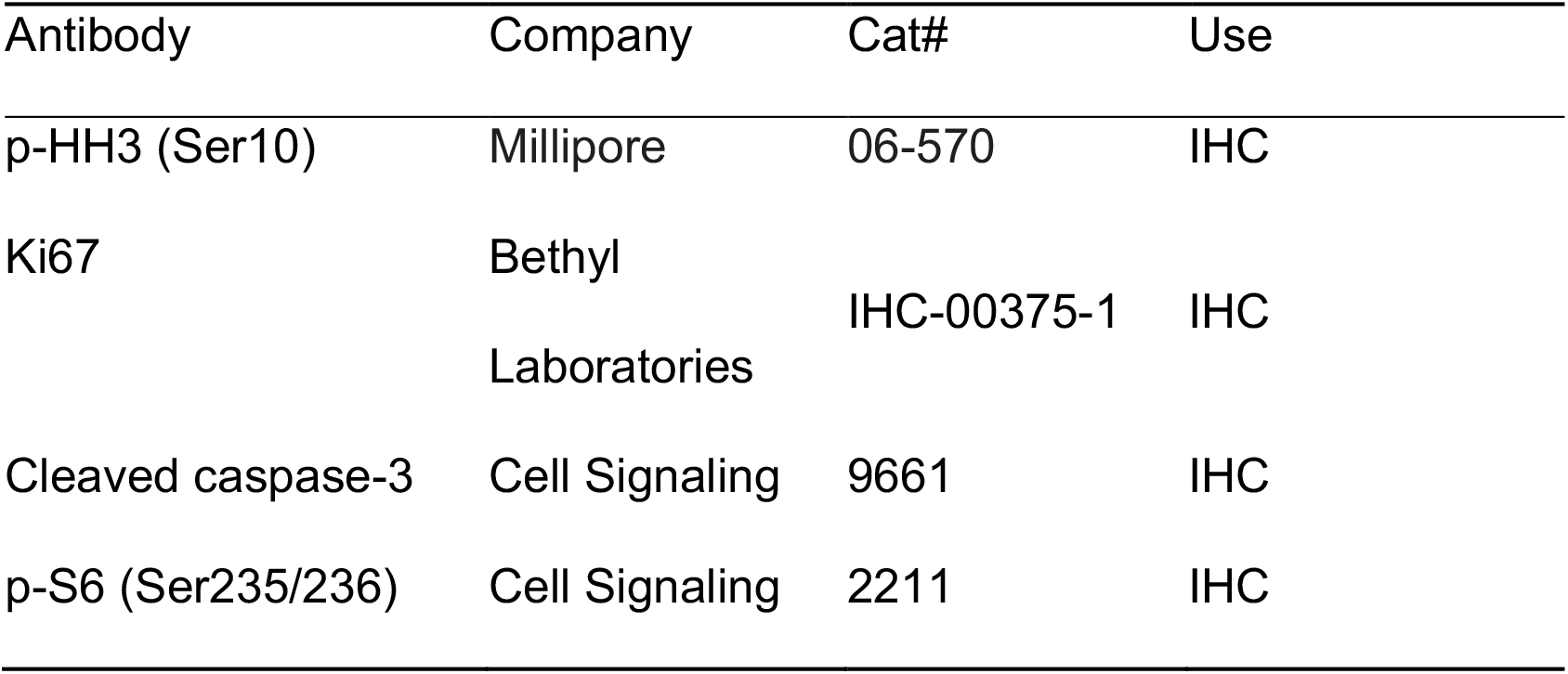
List of antibodies used for this study.

#### Evaluation of metastasis

For all mice harvested at 30 weeks, 10 slides were cut from lungs at 4 µm and every third slide was stained for H&E to total four (slides 1, 4, 7, & 10) H&E-stained slides per lung. H&E-stained slides were then analyzed for metastasis. Metastatic lesions were divided between micrometastasis (area of metastasis is <10.000 µm^2^) and large metastasis (area of metastasis is >10.000 µm^2^). Average metastatic lesions per 4 slides was reported. % coverage of metastasis/lung was analyzed by QuPath and reported.

#### Statistical analyses

Statistical analyses were performed using Microsoft Excel 2016 (Redmond, WA, USA) and GraphPad Prism 9 (San Diego, CA, USA). One-way or two-way ANOVAs and Student’s *t*-tests were used to determine the significance among groups as indicated in the figures or figure legends. Grouped data are presented as mean ± SEM unless otherwise noted. *P* values are indicated in figures or figure legends.

## Results

### Camkk2 deletion initially protects mice from localized disease progression at 15 weeks, but relapses at 30 weeks in a spontaneous prostate cancer mouse model

To understand what role CAMKK2 played in the progression of aggressive prostate cancer, we crossed TRAMP mice with *Camkk2*-/- mice to create TRAMP;*Camkk2*^*-/-*^ (KO) and matched TRAMP;*Camkk2*+/+ (WT) mice. Cohorts of mice were sacrificed at 15 and 30 weeks to study the role of CAMKK2 in prostate cancer initiation and progression, respectively (Figure 1A). Although average prostate (plus potential localized cancer) weights were not significantly different between cohorts (Figure 1B), TRAMP;*Camkk2*^*-/-*^ (KO) mice developed less high-grade PIN at 15 weeks compared to the TRAMP;*Camkk2*^*+/+*^ (WT) cohort (Figure 1C-D). At 15 weeks, KO mice exhibited lower Ki67 staining (Figure 1E), but negligible changes in cleaved caspase 3 (CC3) staining (Supplemental Figure S2A), suggesting decreased proliferation in KO mouse prostates relative to WT controls. Interestingly, at 30 weeks, the KO cohort experienced a relapse (Figure 1C), with no significant difference in Ki67 staining (Figure 1E) or CC3 staining (Supplemental Figure S2B) compared to the WT cohort, suggesting a mechanism of resistance was activated in response to *CAMKK2* loss by 30 weeks.

**Figure 1:**
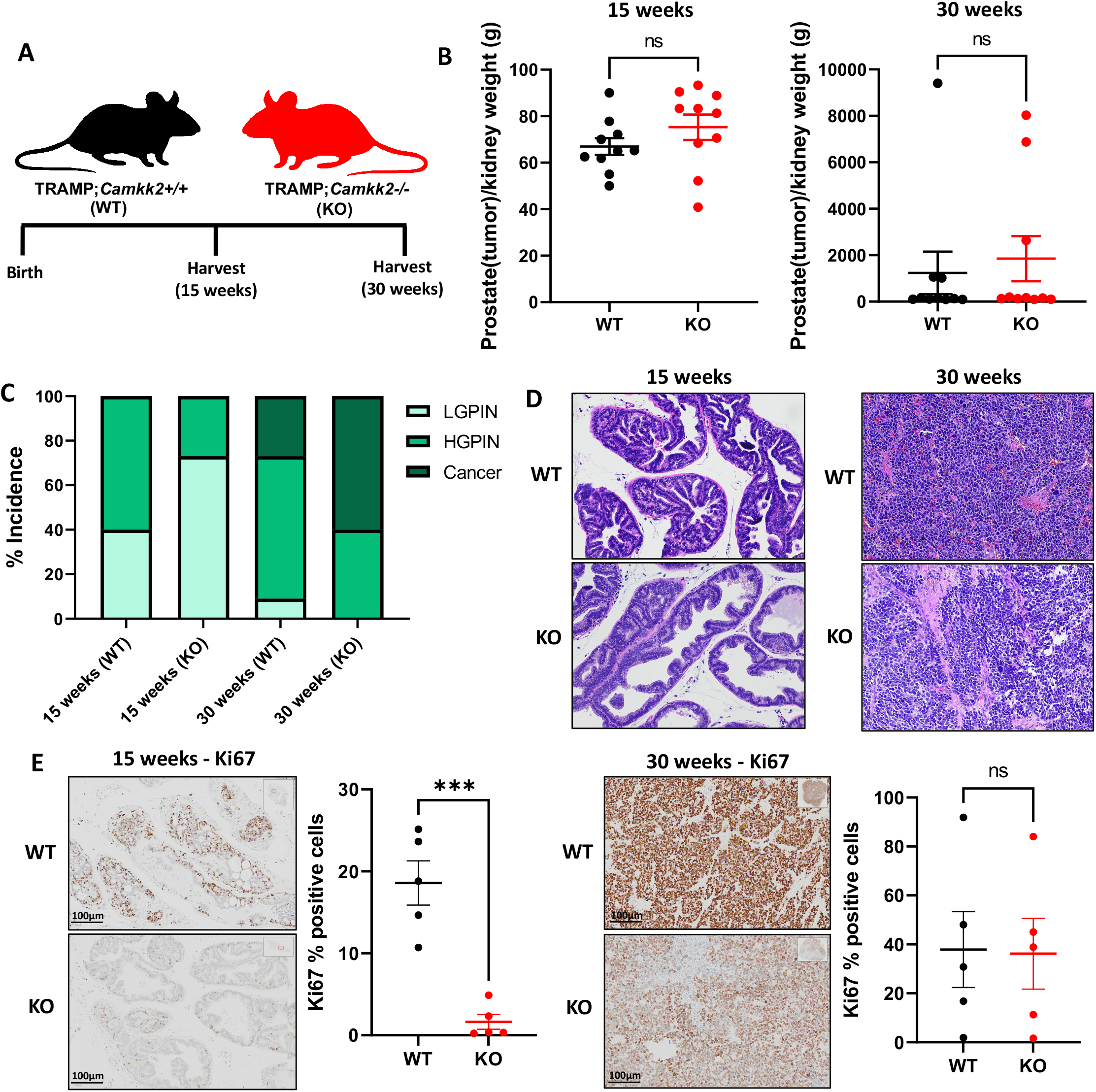
*Camkk2* knockout impairs PIN development at 15 weeks, but disease relapse occurs by 30 weeks in C57BL/6 TRAMP mice. (A) Schema for mouse experiment. (B) Prostate (plus localized cancer if present) weight normalized to kidney weight for 15 weeks (left) (WT n=10, KO n=10) and 30 weeks (right) (WT n=10, KO n=10). (C) % low-grade PIN (LGPIN), high-grad PIN (HGPIN), and cancer incidence among cohorts. (D) Representative images of prostates and tumors. (E) Ki67 staining of tumors at 15 weeks (left) and 30 weeks (right). \*\*\**P* value < 0.001; ns = not significant by *t* test.

### Camkk2 deletion does not change cancer progression rates in castrated TRAMP mice but does alter tumor biology

Next, we castrated mice to determine if CAMKK2 status would impact cancer progression in TRAMP mice under conditions that mimic ADT (Figure 2A). As expected, most prostates atrophied in both WT and KO cohorts of castrated TRAMP mice (Figure 2B), mirroring human prostate response to ADT. Equivalent amounts of prostates/tumors (20%) developed into poorly differentiated/neuroendocrine (PD/NE)-like cancers in both cohorts and prostate/tumor weights as well as Ki67 staining were not significantly different between cohorts (Figures 2B-D; Supplemental Figure S3A) at 30 weeks, indicating that CAMKK2 inhibition did not increase (or decrease) ADT-mediated long-term protection against cancer progression in the context of localized PD/NE tumors. Interestingly, tumors in the KO cohort exhibited much larger regions of necrosis compared to the WT cohort, suggesting differences in tumor biology (Figures 2E and F). Additionally, tumor weight trended lower and CC3 staining trended higher in *Camkk2* KO mice (Figure 2D; Supplemental Figure S3B), suggesting that KO CRPC tumors grew slower and were more necrotic.

**Figure 2:**
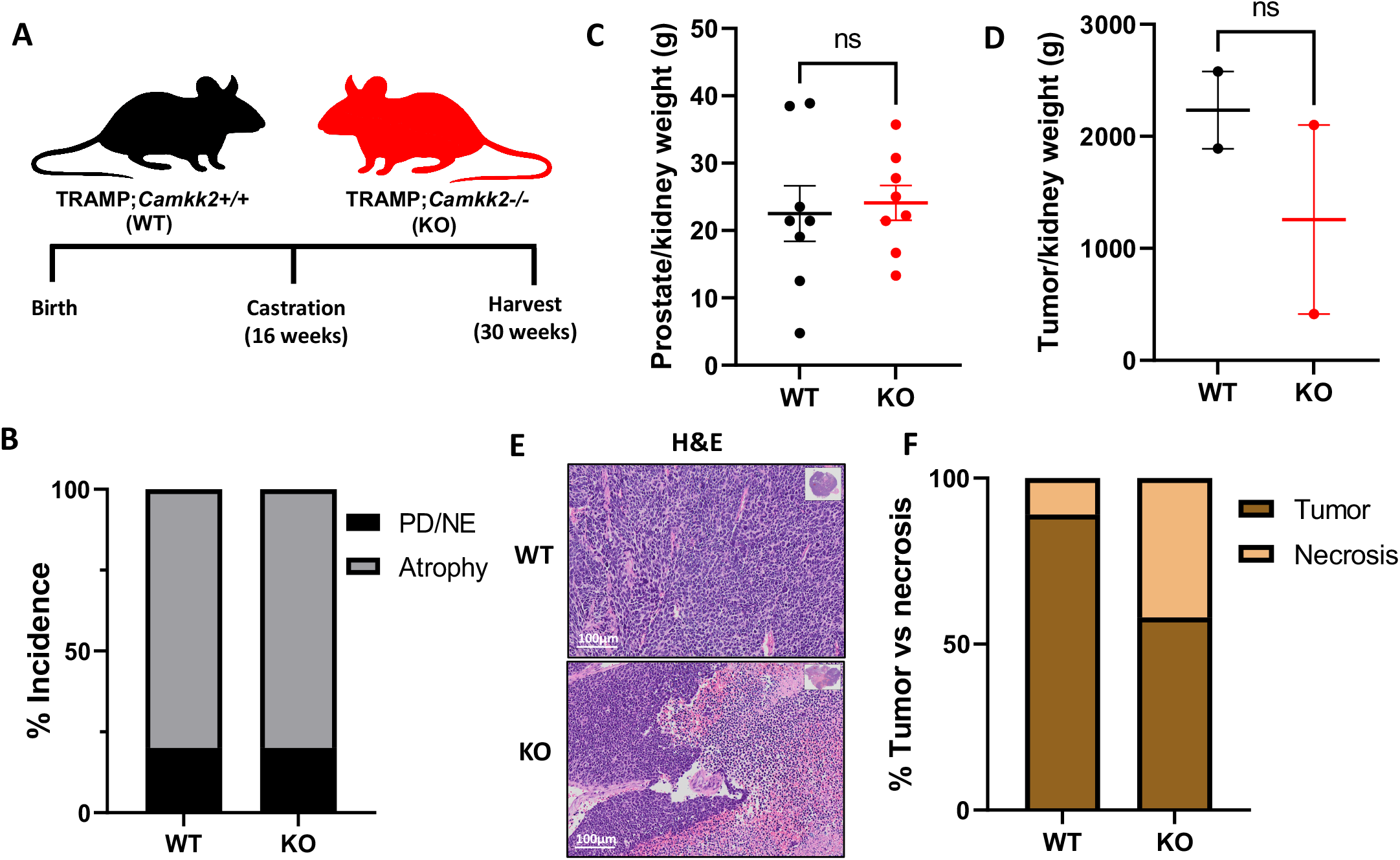
*Camkk2* knockout increases tumor necrosis in castrated TRAMP mice. (A) Schema for mouse experiment. (B) % atrophied prostate and poorly differentiated or neuroendocrine (PD/NE) cancer (WT n=10, KO n=9). (C) prostate (WT n=8, KO n=7) and (D) tumor (WT n=2, KO n=2) weight normalized to kidney weight. (E) Representative images of prostates and tumors. (F) % coverage of tumor cells vs necrosis in tumors (WT n=2, KO n=2) (Fisher’s exact test: *P* < 0.0001). ns = not significant by *t* test.

### Systemic Camkk2 deletion protects against metabolic disorder in a high-fat diet-induced model of obesity and prostate cancer

Previous findings reported that inhibition of CAMKK2 could counteract aspects of metabolic syndrome [19], as well as protect against NAFLD [17]. However, it is unknown whether in prostate cancer models CAMKK2 inhibition would also offset symptoms of metabolic syndrome, a common comorbidity for men with advancing prostate cancer [21]. To test this, we randomized TRAMP mice at 6 weeks to either a high-fat diet (HFD) or normal diet (ND) and collected plasma and organs at 15 and 30 weeks for further analyses (Figure 3A). After 15 weeks on HFD, TRAMP;*Camkk2*+/+ mice displayed fatty livers as assessed by H&E and gross anatomy, which were reduced in the TRAMP;*Camkk2*-/- mice (Figure 3B; Supplementary File 2). However, the protection afforded by *Camkk2* loss against fatty liver in KO mice was only obvious at 15 weeks and was negligible by 30 weeks. Normalized (to kidney) liver weights were not significantly altered by *Camkk2* status (Supplemental Figure S4). Plasma analyses revealed that the TRAMP;*Camkk2*-/- mice had a significant decrease in circulating insulin levels that resulted in reduced HOMA-IR* scores of the non-fasted mice despite maintained blood glucose levels (Figures 3C-E), suggesting improved insulin sensitivity in *Camkk2* KO TRAMP mice. Total cholesterol levels were increased when mice were switched to a HFD but CAMKK2 loss in our model did not significantly change cholesterol levels (Figure 3F). As expected, HFD increased circulating leptin levels in WT TRAMP mice (Figure 3G). *Camkk2* knockout reduced leptin levels in HFD-fed TRAMP mice at 15 but not 30 weeks (Figure 3G), while free fatty acids (FFA) and triglycerides (TG) were reduced at 15 weeks in ND-fed TRAMP mice (Figures 3H and I). Collectively, these metabolic data indicate that CAMKK2 inhibition impacts symptoms of metabolic syndrome including sustained improvements in insulin sensitivity in preclinical models of prostate cancer.

**Figure 3:**
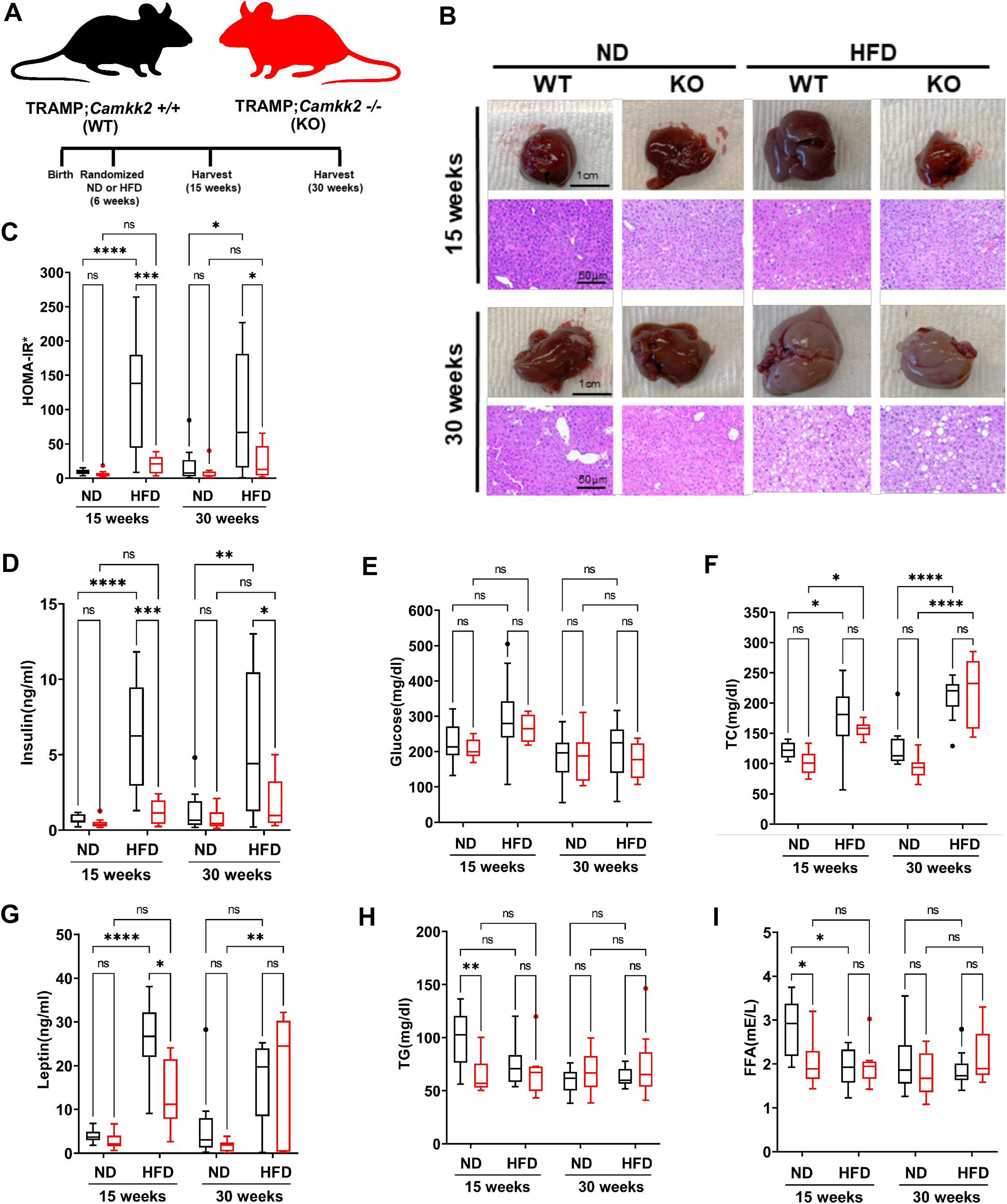
*Camkk2* knockout ameliorates high-fat diet (HFD)-induced insulin resistance. (A) Schema for mouse experiment in which TRAMP;*Camkk2*+/+ (WT) and TRAMP;*Camkk2*-/- (KO) mice were fed either a normal (ND) or high fat diet (HFD) (15 weeks ND: WT n=10, KO n=10; 15 weeks HFD: WT n=10, KO n=8; 30 weeks ND: WT n=10, KO n=9; 30 weeks HFD: WT n=10, KO n=9). (B) Gross morphology and H&E comparison of livers from ND- and HFD-fed TRAMP mice. Raw images and weights of all livers are available in Supplementary File 2. (C-I) Plasma analyses of non-fasted mice. (C) calculated HOMA-IR* scores, and measured levels of (D) insulin, (E) glucose, (F) total cholesterol (TC), (G) leptin, (H) triglycerides (TG), and (I) free fatty acids (FFA). \**P* < 0.05, \*\*\**P* < 0.0001; Two-way ANOVAs.

### Camkk2 deletion impairs the metastatic colonization of NEPC tumors

Using the TRAMP model allowed us to investigate the impact of a systemic *Camkk2* knockout, not just on spontaneously developed localized prostate cancer, but also the subsequent metastasis to distant organs such as the lung. We observed that *Camkk2* knockout, regardless of diet, reduced localized prostate cancer at 15 weeks but not at 30 weeks (Figures 4A-B). *Camkk2* status did not influence prostate/primary tumor weight in obese TRAMP mice (Supplemental Figure S5). Notably, a HFD, while having minimal impact on the overall incidence of metastasis, promoted metastatic colonization relative to mice maintained on a normal diet (Figures 4C-G, Supplementary File 3). However, this HFD-mediated metastatic colonization was dependent on CAMKK2 (Figures 4F-G; Supplementary File 3). In fact, metastatic lesions were clearly visible to the naked eye in the majority of the HFD-fed TRAMP;*Camkk2*+/+, but only one of the TRAMP;*Camkk2*-/- histology slides. Although not significant, there was a trend towards decreased overall incidence of metastasis in both ND- and HFD-fed TRAMP mice when *Camkk2* was ablated (Figure 4E). Importantly, the metastases observed in this model were all NEPC. Taken together, CAMKK2 inhibition suppressed the metastatic colonization of NEPC tumors in a HFD-driven GEMM of aggressive prostate cancer.

**Figure 4:**
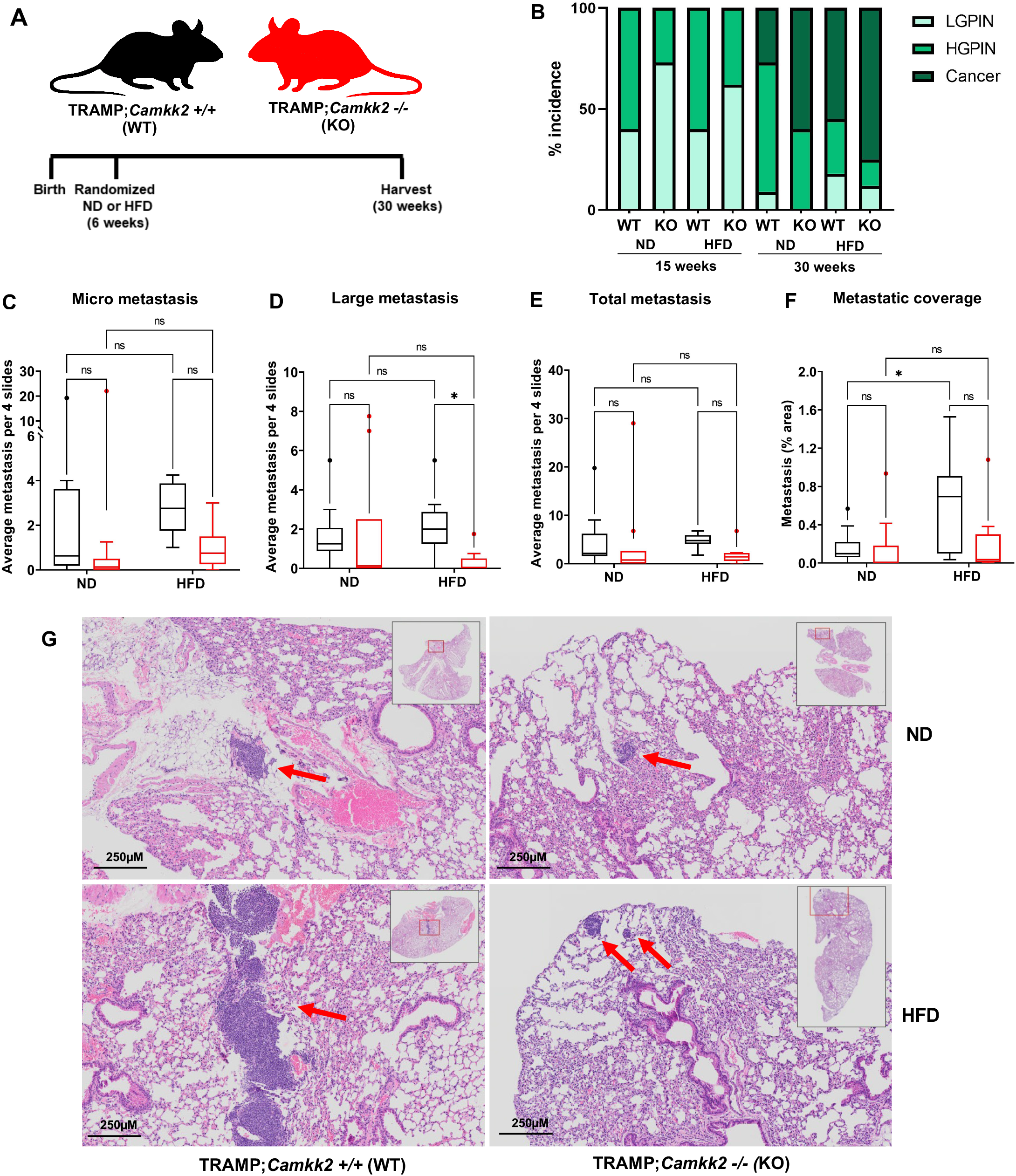
*Camkk2* knockout decreases high-fat diet-induced metastatic colonization of NEPC in TRAMP mice. (A) Schema for mouse experiment in which C57BL/6 TRAMP;*Camkk2*+/+ and TRAMP;*Camkk2*-/- mice were fed either a high-fat diet (HFD) or normal diet (ND) (HFD WT n=8; HFD KO n=8; ND WT n=9; ND KO n=10). (B) % low-grade PIN (LGPIN), high-grad PIN (HGPIN), and cancer incidence among cohorts. Lung metastasis was quantified by counting incidents of (C) micrometastases (area of metastasis is < 10.000 µm^2^), (D) large metastases (area of metastasis is > 10.000 µm^2^), or (E) any metastatic lesions. In addition, (F) image analysis was performed on lung H&E slides to quantify the % metastatic area relative to total lung (total lung = 100%). (G) Representative H&E stains of lung metastases found. For full data set and analyses, see Supplementary File 3. \**P* < 0.05; Two-way ANOVA.

### Host Camkk2 ablation decreases cancer cell size and mTOR signaling in the TRAMP GEMM model of prostate cancer

When analyzing cancer cell morphology between cohorts in the TRAMP mice, we observed a subtle, yet highly reproducible (*P* < 0.0001; n > 1 × 10^6^ cells/group) decrease in cell size (area and predicted volume) in both ND and HFD KO cohorts compared to WT mice (Figures 5A-B). Given the lower circulating insulin levels in *Camkk2* KO mice (Figure 3D), we hypothesized that this change in cell size could be due to decreased mTOR activity, a downstream effector of insulin signaling and major regulator of cell size [26-28]. To test this, we stained tumors for p-S6, a marker of mTOR activation. As expected, increased p-S6 staining was observed in both primary tumors and metastatic lesions in HFD-fed mice relative to control-fed mice (Figures 5C). Interestingly, p-S6 levels were significantly decreased only in the metastatic lesions from HFD-fed TRAMP mice lacking *Camkk2* relative to WT (Figure 5C), mirroring the effects we observed on metastatic colonization (Figures 4F-G). While there was a trend toward decreased p-S6 in primary tumors, the change in p-S6 did not reach significance.

**Figure 5:**
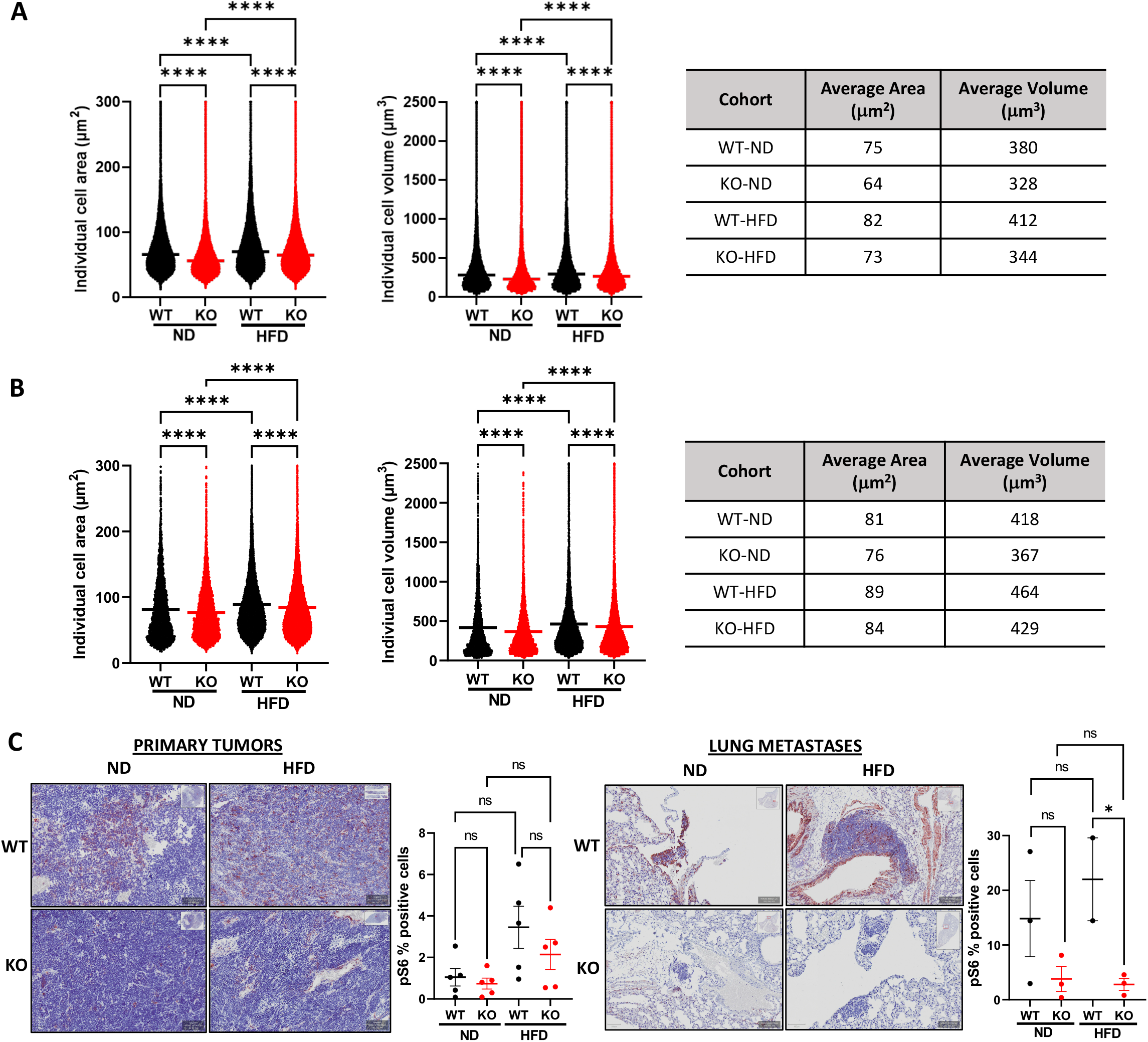
Host *Camkk2* ablation decreased cancer cell size and mTOR signaling in TRAMP mice. (A) Individual cell areas and predicted cell volumes for primary tumors derived from 30-week old TRAMP;*Camkk2+/+* and TRAMP;*Camkk2-/-* mice described in Figure 4A. (B) Individual cell areas and predicted cell volumes for lung metastases derived from the same TRAMP;*Camkk2+/+* and TRAMP;*Camkk2-/-* mice. (C) Representative images (*left*) and quantification (*right*) of p-S6 IHC stains (n=5/group) from tumors in (A) and lung metastases in (B). ns = not significant; \**P* < 0.05, \*\*\*\**P* < 0.0001. For each morphometric analysis, n > 1 × 10^6^ cells/group.

### Host Camkk2 ablation decreases tumor growth and cancer cell size in syngeneic mouse models of prostate cancer: Evidence of cancer cell-extrinsic roles for CAMKK2 in prostate cancer

To separate CAMKK2’s tumor extrinsic functions from its intrinsic functions in cancer, we subcutaneously injected cells from two syngeneic, *Camkk2*-intact/WT mouse prostate cancer lines (RM-9 and TRAMP-C2) into litter-matched *Camkk2*+/+ (WT) or *Camkk2*-/- (KO) host mice (Figure 6A). While TRAMP-C2 is hormone-sensitive, RM-9 exhibits *de novo* resistance to androgen deprivation *in vitro* and *in vivo* (Supplemental Figure S6). Both syngeneic tumors grew more slowly in the KO mice indicating that cancer cell-extrinsic CAMKK2 promotes prostate cancer progression (Figures 6B-C; Supplemental Figure S7). Like observed in the TRAMP mice, both syngeneic tumors exhibited smaller average cell sizes when propagated in *Camkk2* KO host mice compared to WT mice (Figures 6D-E). Also consistent with what we observed in TRAMP metastatic lesions, tumors grown in mice lacking *Camkk2* exhibited decreased mTOR activity as assessed by p-S6 IHC (Figures 6D-E). Together, our data suggest that, in addition to the previously described cancer cell-intrinsic role of CAMKK2 in prostate cancer [3-5,29,30], CAMKK2 also promotes prostate cancer progression via tumor extrinsic mechanisms. Specifically, we propose that CAMKK2 enables metabolic syndrome, which causes increased levels of circulating insulin or some insulin-like molecule that can promote oncogenic mTOR signaling in distant prostate cancer cells (Figure 7).

**Figure 6:**
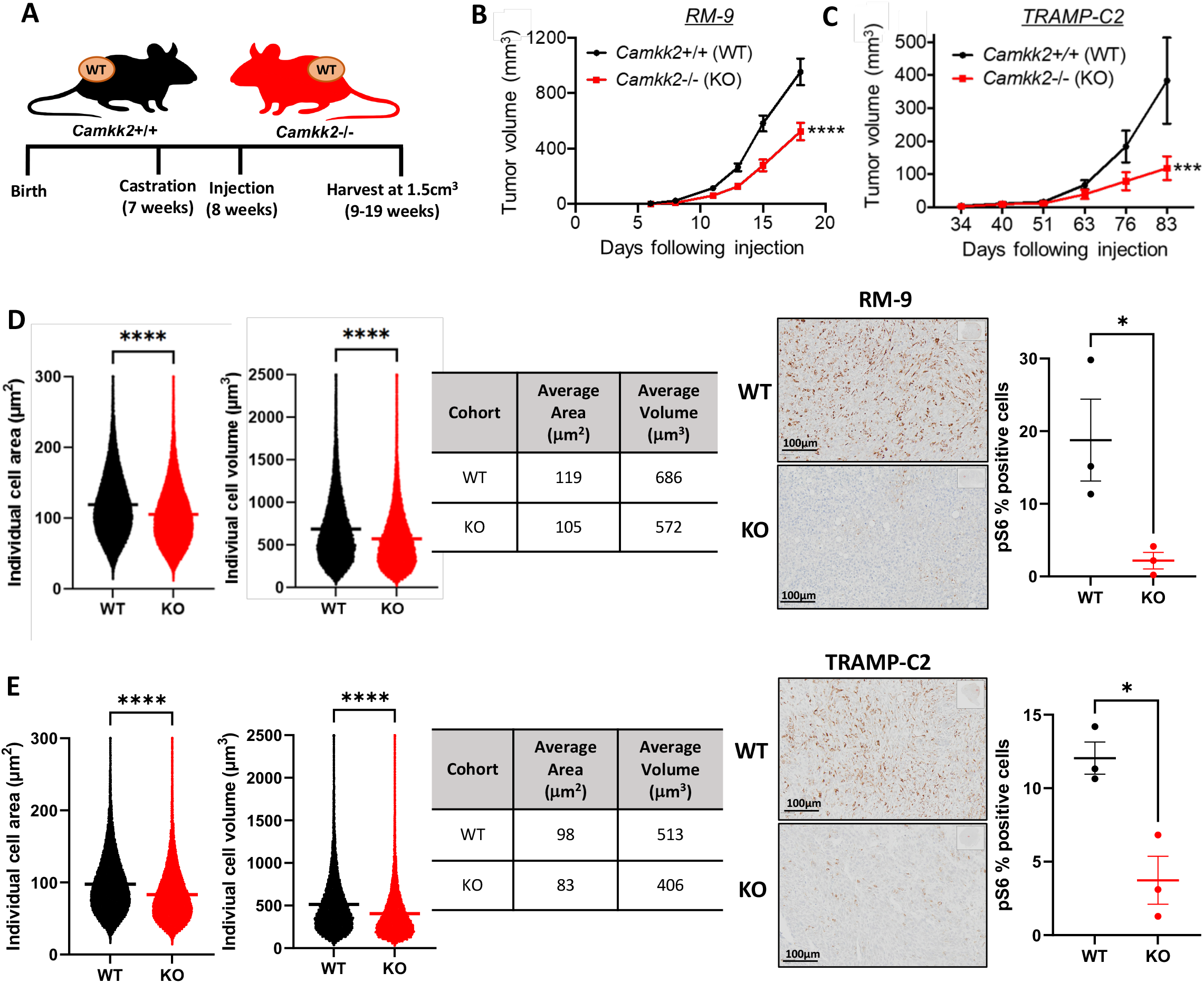
Host *Camkk2* ablation decreased tumor growth, cancer cell size, and mTOR signaling in syngeneic mouse models of prostate cancer. (A) Schema for syngeneic mouse experiment. C57BL6 RM-9 (B) and TRAMP-C2 (C) murine prostate cancer cells were subcutaneously injected into syngeneic *Camkk2*+/+ and *Camkk2*-/- host mice and tumor growth was monitored over time. (D) RM-9 and (E) TRAMP-C2 average cell areas and average cell volumes were quantified from harvested tumors (*left*). In addition, % positive staining of p-S6 in syngeneic tumors (WT n=3, KO n=3) was quantified (*right*). ns = not significant; \**P* < 0.05, \*\*\*\**P* < 0.0001. For each morphometric analysis, n > 1 × 10^6^ cells/group.

**Figure 7:**
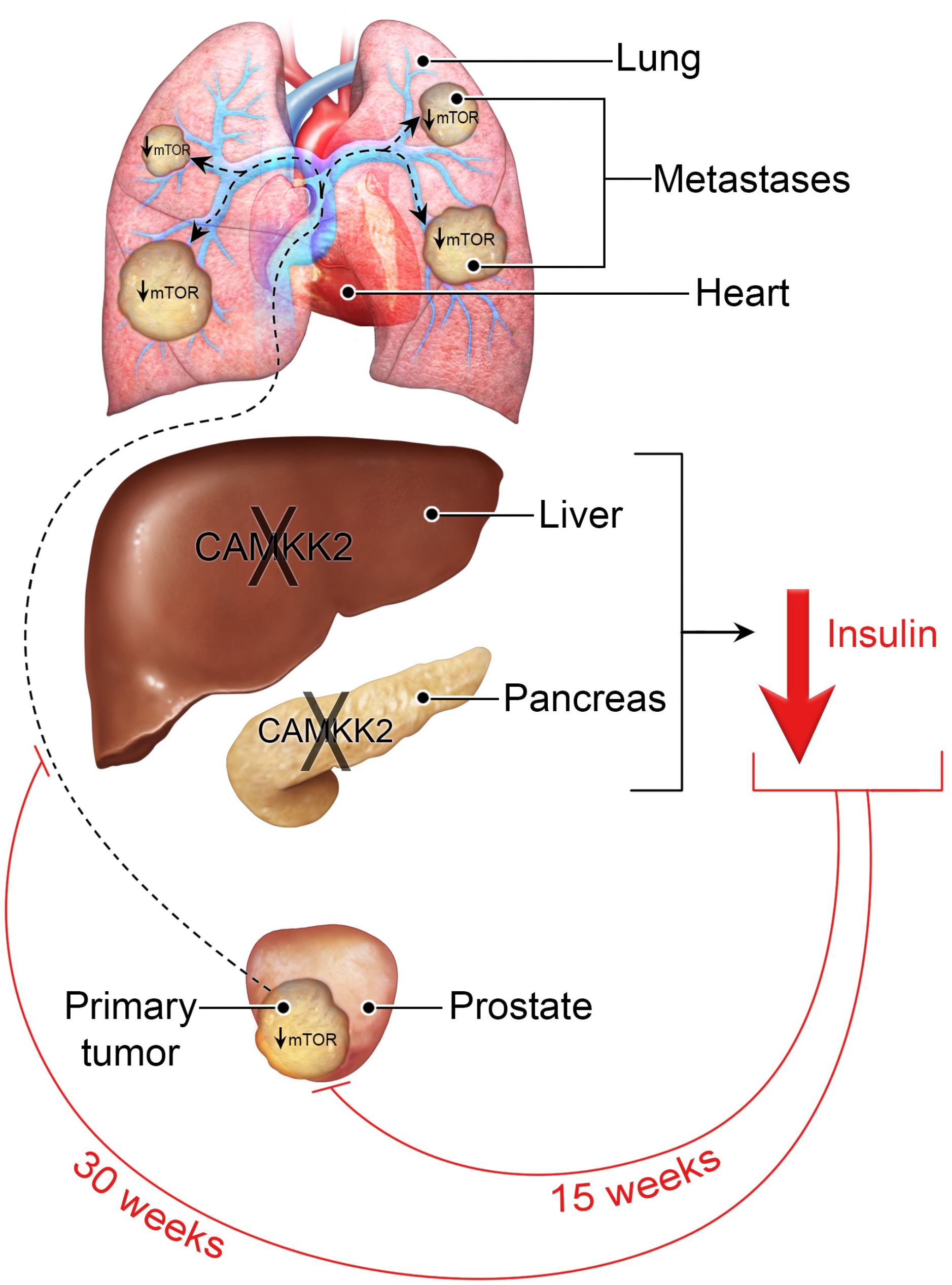
Working model for CAMKK2’s proposed tumor-extrinsic role in prostate cancer. We propose that *Camkk2* KO in the liver and/or pancreas protects against metabolic syndrome. One consequence of this is lowered circulating insulin (or some insulin-like molecule), which slows early primary tumor growth and later metastatic colonization through decreasing cancer cell-intrinsic, oncogenic mTOR signaling.

## Discussion

Although the initial report of CAMKK2’s functional role in prostate cancer noted effects on prostate cancer cell migration and invasion [3], to our knowledge, CAMKK2’s role in prostate cancer metastasis *in vivo* had not been described until this study. Interestingly, we also discovered an oncogenic role for CAMKK2 in driving the progression of NEPC, an extremely aggressive subtype of the disease [31]. This was surprising given that *CAMKK2* is a direct target of AR in both hormone-sensitive and CRPC states [3,4,32]. Because of this direct regulation by AR, CAMKK2’s functions in prostate cancer were largely assumed to be limited to classic, AR+ adenocarcinomas. However, prior reports suggesting roles for CAMKK2 in an AR-cell line in culture [33] and here *in vivo* indicate roles for CAMKK2 even in AR-indifferent prostate cancers. Given the lack of effective treatments for this highly aggressive cancer subtype, it suggests that the use of CAMKK2 inhibitors, which are currently in development [24,34-38], warrant further investigation for the treatment of NEPC.

While CAMKK2 is an interesting therapeutic target due to its role in prostate cancer cell biology, prior reports indicate that CAMKK2 inhibitors may have additional benefits for prostate cancer patients [20]. One of the most common comorbidities for men undergoing ADT is metabolic syndrome [21]. This side effect of AR-targeting treatment may contribute in part to the limited improvements in overall survival observed in patients since ∼2012 despite the development and FDA approval of several new prostate cancer therapies [1]. Notably, we observed simultaneous anti-cancer effects with improvements in liver pathology and insulin sensitivity in our HFD/obesity-driven TRAMP model following *Camkk2* ablation (Figure 3). These data indicate that systemic CAMKK2 inhibitors may provide dual benefits for men with advanced prostate cancer, inhibition of both the cancer and linked comorbidities. Consistent with the concept that selective CAMKK2 inhibitors could be well tolerated in patients, mice harboring germline *Camkk2* deletions are viable, fertile, and have long lifespans [22]. Clearly, the observed benefits of CAMKK2 inhibition on systemic metabolism will need to be formally tested in diverse models of ADT-driven metabolic syndrome to provide a more comprehensive evaluation of anti-CAMKK2-mediated efficacy.

To date, CAMKK2’s oncogenic effects in prostate cancer have solely been attributed to its known cancer cell-autonomous functions [3-5,29,30,33]. Given the multiple effects of *Camkk2* ablation on cancer and macrometabolism, perhaps not surprisingly, here we observed for the first time an additional cancer cell-extrinsic role for CAMKK2 in prostate cancer progression (Figure 6). It is our working hypothesis that CAMKK2 activity in peripheral metabolic organs such as the liver and pancreas as previously reported [18,19] facilitate higher circulating levels of insulin or some insulin-like molecules (ex. IGF-1), known mitogens, to restore or maintain metabolic homeostasis (Figure 7). A byproduct of the elevated insulin is increased oncogenic mTOR signaling at distant sites. These data are consistent with a growing body of literature that suggests that increases in circulating levels of insulin or insulin-like molecules may be more directly promoting the progression of multiple cancer types more so than other conditions associated with obesity/high body mass index (BMI) [39,40]. To that end, prior studies have demonstrated that insulin and IGF-1 can directly stimulate the proliferation of a subset of cancer cells including TRAMP-C3 prostate cancer cells [41,42]. Arguing against a specific role for IGF-1 is a prior report that liver-specific deletion of *Igf1* in TRAMP mice did not impact tumor progression [43]. Likewise, we also observed minimal effects of *Camkk2* ablation on metastasis in lean mice. Whether liver-specific deletion of *Igf1* in the context of obesity would alter disease outcome is unknown. While our data suggest that CAMKK2 functions in tissues or organs that control systemic metabolism, we do not yet know the exact tissue(s) through which CAMKK2 is functioning. Based on prior reports studying CAMKK2 in the liver and pancreas [18,19], we speculate that CAMKK2 activity in one or both organs may play a significant role in prostate cancer progression.

Previously, a different cancer cell-extrinsic role was reported in breast cancer involving the regulation of the anti-tumor immune response through CAMKK2’s functions in macrophages [44] and more recently in a preclinical model of lymphoma, myeloid-derived suppressor cells (MDSCs) [45]. While our initial histological analysis did not note any overt changes in tumor immune cell infiltration, more in-depth profiling is required to determine if an immunosuppressive switch is occurring within the prostate cancer tumor microenvironment. Moreover, established links between obesity and inflammation indicate that it will be important to determine: 1) if there is crosstalk between these two areas, 2) what role CAMKK2 may play in their regulation, and 3) the ultimate impact on cancer.

Systemic *Camkk2* ablation slowed disease progression in the TRAMP GEMM of prostate cancer (Figure 1) similar to what was observed in the Pb-Cre4;*Pten*^f/f^ GEMM of prostate cancer [9]. Unlike what was observed in Pb-Cre4;*Pten*^f/f^ mice, at 30 weeks TRAMP;*Camkk2*-/- mice exhibited relapsed local prostate cancer progression. The exact cause of this difference is unclear, but we speculate that it is due in part to TRAMP mice being an inherently more aggressive model of prostate cancer compared to Pb-Cre4;*Pten*^f/f^ [46]. The disease relapse observed in the TRAMP mice indicates some type of acquired resistance to CAMKK2 inhibition that could be important when designing future combination therapies should CAMKK2 inhibitors make it to the clinic. The mechanism of resistance may coincide with the onset of NEPC lesions, which are known to occur in TRAMP (albeit less frequently on the C57BL/6 genetic background used in this study compared to the more commonly used C57BL/6;FVB mixed background), but not Pb-Cre4;Pten^f/f^ mice, which maintain an adenocarcinoma phenotype [46]. While pathogenesis in Pb-Cre4;*Pten*^f/f^ mice is driven by the loss of a single tumor suppressor (PTEN), the TRAMP model is driven by the inactivation of multiple tumor suppressors (RB1, p53, and to a lesser extent PP2A). Although loss of function of all these tumor suppressors is common in advanced prostate cancer [47], deletion or mutation of any two of the combination of *PTEN, RB1, TP53* predicts poor response to AR-targeted therapy and decreased overall survival [48]. As such, these genetically defined prostate cancers have been classified as aggressive variant prostate cancers (AVPCs), of which NEPCs are typically included [31]. Curiously, while disease relapse was observed at 30 weeks in TRAMP mice at the primary site (prostate) regardless of diet, systemic *Camkk2* loss dramatically impaired the metastatic colonization of the lungs at the same timepoint in HFD-fed TRAMP mice (Figure 4). The differences in tumor growth at these two sites (primary and metastasis) mirrored changes in p-S6 staining, indicating that mTOR signaling was reactivated in the primary tumor but continued to be suppressed in the lung macrometastases. These data suggest that tumor-extrinsic CAMKK2 signaling may have a greater role in metastasis compared to primary tumor growth.

It is unknown if the tumor-extrinsic effects we observed for CAMKK2 are specific for any genetic subtypes. Neither the GEMM (TRAMP) or syngeneic mouse models (RM-9 and TRAMP-C2) used in this study are driven by alterations in PI3K-AKT signaling, unlike the Pb-Cre4;*Pten*^f/f^ GEMM. We speculate that the higher levels of circulating insulin in the *Camkk2* WT mice, exacerbated during obesity, promote aberrant PI3K-AKT-mTOR signaling that potentially provides the missing third oncogenic piece to AVPCs previously driven by the other two genetic hallmarks of AVPC, *RB1* and *TP53* genetic alterations. This would be consistent with GEMM studies demonstrating that tumors harboring alterations in all three tumor suppressors (*Pten, Rb1, Tp53*) have *de novo* resistance to AR-targeted therapy whereas the presence of only two of the three alterations requires an additional period of acquired resistance [49,50]. Tumors in Pb-Cre4;*Pten*^f/f^ mice may be less sensitive to insulin/tumor-extrinsic CAMKK2 functions since there is already high basal mTOR signaling. However, recent work has suggested that HFD-mediated hyperinsulinemia may also accelerate disease progression in Pb-Cre4;*Pten*^f/f^ (FVB) mice [51]. Future studies directly comparing these models will help to resolve this issue.

CAMKK2-regulated differences in cancer cell size and mTOR signaling (assessed by p-S6 IHC) combined with the sustained alterations in circulating insulin levels suggest that insulin mediates CAMKK2’s effects on cancer via control of systemic metabolism. However, we cannot rule out contributions from other factors linked to metabolic syndrome such as leptin, triglycerides and free fatty acids that were altered by CAMKK2 deficiency at 15 weeks but not 30 weeks (Figure 3). For example, insulin may have a substantial role in metastasis, whereas additional factors like leptin, triglycerides and free fatty acids could have important roles in promoting localized, primary tumor growth (consistent with the observed prostate pathology at 15 and 30 weeks (Figures 1 and 4)). Indeed, there is compelling preclinical and clinical data supporting important roles for local factors such as fats and paracrine growth factors in promoting primary tumorigenesis [52-54]. Why CAMKK2 inhibition does not sustain decreases in these local factors is unclear but it may be linked to its regulation of adipocyte differentiation, which could change the cellular milieu over time [55]. Our data suggest that treatment of primary prostate cancer may necessitate that CAMKK2 inhibitors be combined with additional therapies to overcome the localized resistance mechanisms.

Efforts are ongoing by multiple groups to develop new potent and selective CAMKK2 inhibitors with the goal to create a clinical-grade drug for use in patient trials [24,34-38]. The data presented here indicate that CAMKK2 inhibitors may have additional benefits for patients beyond what was initially anticipated. First, our data suggest that CAMKK2 inhibitors could have value in the treatment of distant metastases and/or NEPC, leading causes of prostate cancer mortality. In addition, CAMKK2 inhibitors may offer patients a “2 for 1” deal by treating not only the cancer directly, but also disease-linked comorbidities like metabolic syndrome. Moreover, if CAMKK2’s hepatic functions are proven to be a major driver of prostate cancer progression, this could present new opportunities from a drug development/pharmacokinetic standpoint, given that many drugs often accumulate in the liver [56].

### Caveats

Although the data presented here provide compelling new evidence for CAMKK2 functions in prostate cancer progression and potentially related effects on systemic metabolism, we are cognizant that this initial study has several caveats. First, we are yet to functionally link the CAMKK2-mediated changes in circulating insulin levels to the observed changes in cancer cell size, mTOR signaling, and cancer progression. Experiments testing the necessity and sufficiency of insulin or insulin-like molecules will be essential to solidify our working model (Figure 7). Second, we did not measure total body weights. In future studies, this would be an important parameter to measure since *Camkk2*-/- mice were initially described to have an altered appetite [22]. However, CAMKK2’s impact on circulating insulin levels was later shown to be independent from its effects on appetite [19]. Third, our plasma measurements were not done on fasted mice because we did not want to add additional variables when assessing tumor biology. Future studies will require *bona fide* measures of glucose homeostasis such as fasting insulin and glucose levels, as well as glucose and insulin tolerance tests. Fourth, we did not have a HFD-fed, castrated TRAMP cohort. In prostate cancer patients, ADT is known to increase the risk of metabolic syndrome [20,21,57]. We would predict that castrated mice fed a HFD-diet may benefit even more from CAMKK2 inhibition, a scenario that remains to be tested. Fifth, we did not use a HFD with either of our syngeneic mouse models, nor did we collect blood for plasma analysis. Hence, we had to rely on historical measurements [17,18] as well as comparisons to our TRAMP experiments. Based on prior observations, we anticipate *Camkk2* ablation could have even more pronounced effects in obese syngeneic mouse models. Finally, although we focused on mTOR signaling due to the changes in insulin, we cannot rule out a role for the other predominant growth-control pathway, the Hippo signaling cascade [58].

## Conclusions

In summary, our data demonstrate new oncogenic roles for CAMKK2 in metastatic colonization and NEPC progression. In addition, genetic targeting of *Camkk2* in immune-intact mouse models of obesity-driven prostate cancer demonstrated simultaneous anti-cancer efficacy and improvements in whole body metabolic homeostasis that we propose may be linked via a novel tumor-extrinsic, CAMKK2-mediated mechanism of action. Together, these data force us to reevaluate the potential mechanisms of action from prior data reporting anti-cancer effects in models leveraging systemic CAMKK2 inhibition via either germline genetic alterations (ex. *Camkk2*-/- mice) or small molecule inhibitors (ex. STO-609). Nevertheless, the multiple mechanisms of action of CAMKK2 underscore the potential of CAMKK2 inhibitors for the treatment of cancer as well as other diseases such as diabetes, fatty liver disease, and more.

## Funding

This work was supported by grants from the National Institutes of Health (NIH R01CA184208 and P50CA140388 (D.E.F.)), American Cancer Society (RSG-16-084-01-TBE (D.E.F.), and a grant from the Mike Slive Foundation for Prostate Cancer Research (D.E.F.). J.W.S. and J.S.O. are funded by National Health and Medical Research Council (NHMRC) grants GNT1138102 and GNT1145265, respectively. This work was also supported by an American Legion Auxiliary Fellowship (D.A.). Some of the histology was performed with the CCSG-funded MDACC Research Histology Core Laboratory, NIH grant P30CA016672. This project was also supported by the Mouse Metabolism and Phenotyping Core at Baylor College of Medicine with funding from the NIH (UM1HG006348, R01DK114356, R01HL130249).

## Institutional Review Board Statement

The animal study protocol was approved by the Institutional Review Board of the University of Texas MD Anderson Cancer Center (IACUC protocol#: 00001738-RN01, approved 10/4/2021) and the University of Houston (IACUC protocol#: 14-039, approved 4/17/2017).

## Acknowledgments

We would like to thank Kelly Kage (UT MD Anderson Cancer Center) for artistic assistance preparing Figure 7, Pratip Saha and the Baylor College of Medicine (BCM) Mouse Metabolic Phenotyping Core for assisting with measurements of various blood parameters, Joseph Daniele (UT MD Anderson Cancer Center) for help with image analysis, Brian York and Anthony Means (BCM) for helpful conversations and providing the original C57BL/6 *Camkk2*-/- and *Camkk2* floxed mice, David Stewart (University of Houston) for providing the Sox2-Cre mice, and Raghu Kalluri (UT MD Anderson Cancer Center), Sanghyuk Chung (University of Houston) and Barbara Foster (Roswell Park Comprehensive Cancer Center) for advice regarding the analysis of metastatic lesions, histology, and use of the TRAMP mice.

## Conflicts of Interest

D.E.F. has received research funding from GTx, Inc and has familial relationships with Hummingbird Bioscience, Maia Biotechnology, Alms Therapeutics, Hinova Pharmaceuticals and Barricade Therapeutics. The other authors report no potential conflicts of interest. The funders had no role in the conceptualization of the study or writing of the manuscript, or in the decision to publish this article.

