## Supplemental File 1 for "Systemic ablation of *Camkk2* impairs metastatic colonization and improves insulin sensitivity in TRAMP mice: Evidence for cancer cell-extrinsic CAMKK2 functions in prostate cancer"

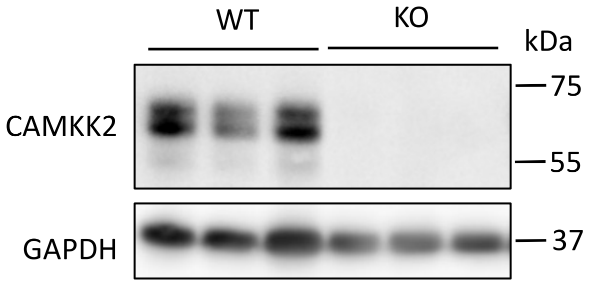


**Supplemental Figure S1: Germline *Camkk2* knockout in C57BL/6 mice confirmed by western blot analysis of brain lysates from *Camkk2*+/+ (WT) and *Camkk2*-/- (KO) mice.**

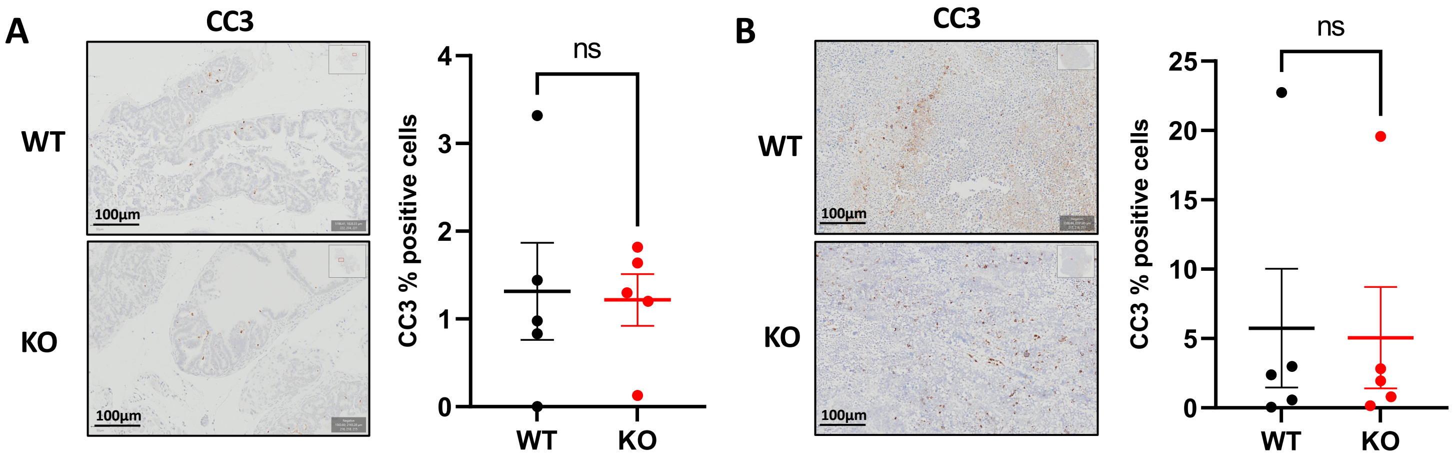


**Supplemental Figure S2: *Camkk2* knockout did not affect apoptosis as measured by cleaved caspase-3 (CC3) staining in prostates/tumors from TRAMP mice.** Cleaved caspase-3 (CC3) staining of prostate/tumors at (A) 15 weeks (WT n=5, KO n=5) and (B) 30 weeks (WT n=5, KO n=5) shows no significant change in staining. ns = not significant by *t* test.

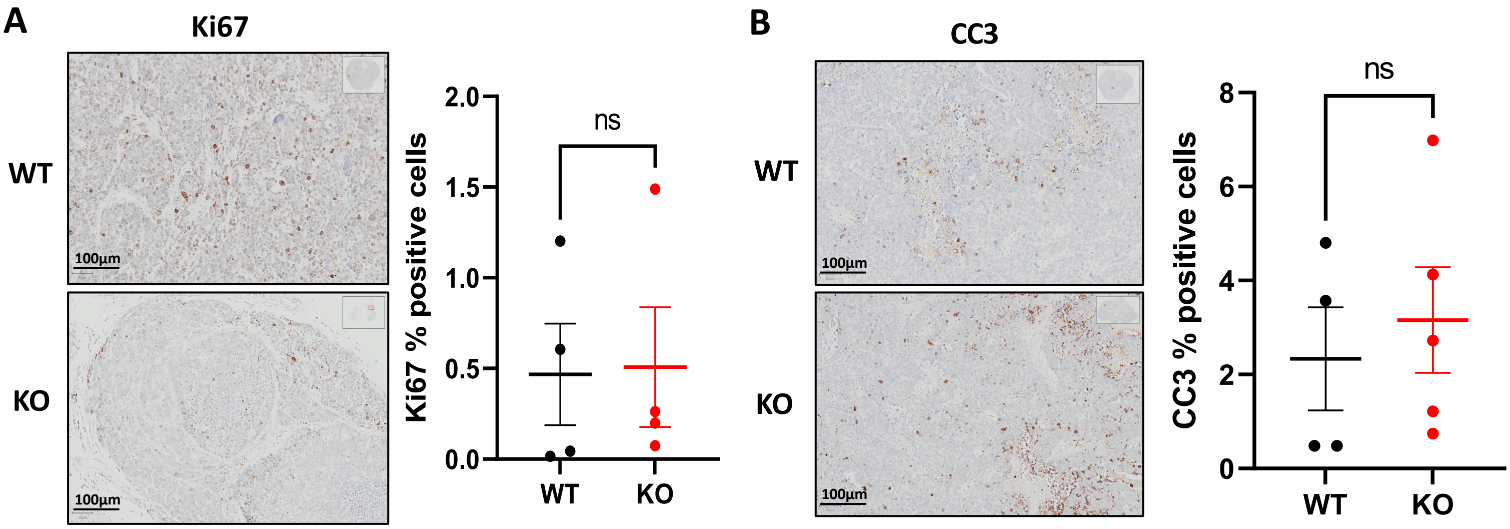


**Supplemental Figure S3: Effects of *Camkk2* knockout in castrated TRAMP mice on proliferation and apoptosis.** (A) Ki67 (WT n=4, KO n=4) and (B) cleaved caspase-3 (CC3) IHC analyses of castrated TRAMP mice (WT n=4, KO n=5). ns = not significant by *t* test.

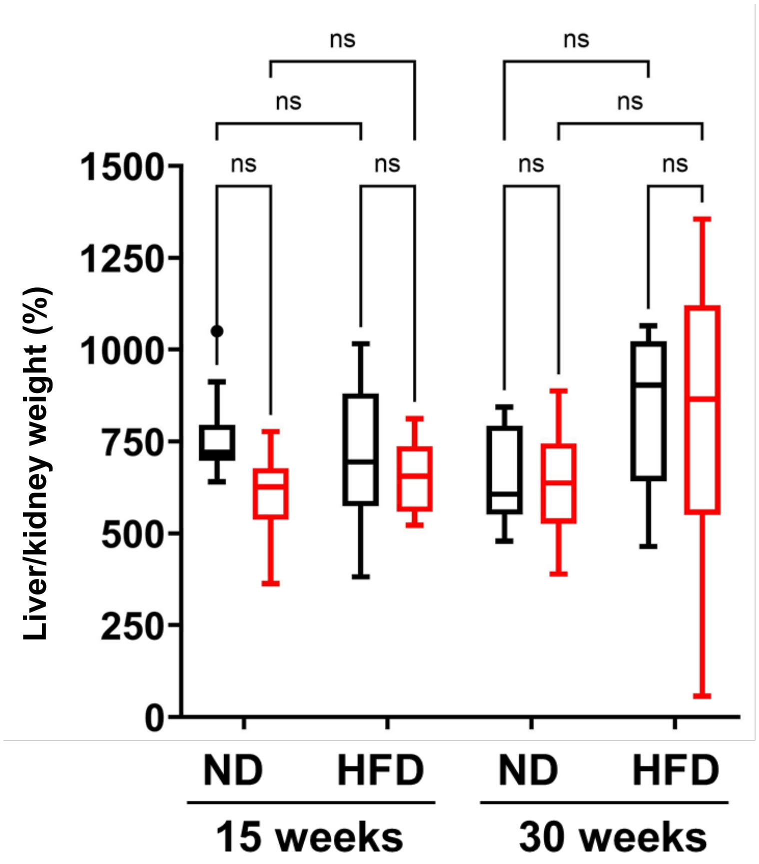


**Supplemental Figure S4: CAMKK2 status did not influence liver weights in TRAMP mice.** Livers were weighed and normalized to kidney weights in 15- and 30-week old TRAMP;*Camkk2*+/+ (black) and TRAMP;*Camkk2*-/- (red) mice fed normal (ND) or high-fat diet (HFD) (15 weeks ND: WT n=10, KO n=10; 15 weeks HFD: WT n=10, KO n=8; 30 weeks ND: WT n=10, KO n=9; 30 weeks HFD: WT n=10, KO n=9). ns = no significance; Two-way ANOVA.

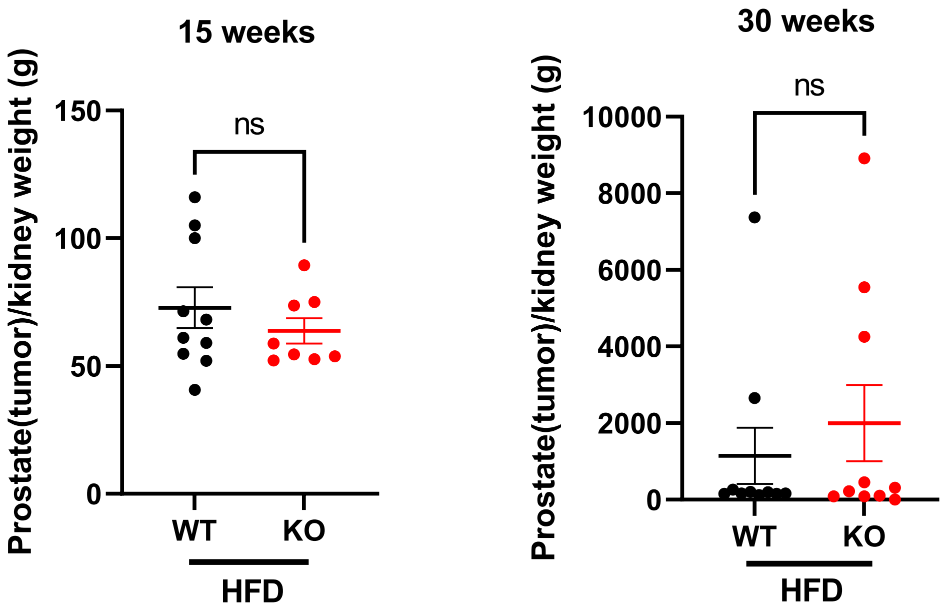


**Supplemental Figure S5: *Camkk2* knockout does not impact prostate/primary tumor weights in high-fat diet (HFD)-fed mice.** Prostate/primary tumor weights normalized to kidney weight at 15 (left) or 30 (right) weeks in HFD-fed mice (15-week HFD WT n=10; 15-week HFD KO n=8; 30-week HFD WT n=11; 30-week HFD KO n=10). ns = no significance.

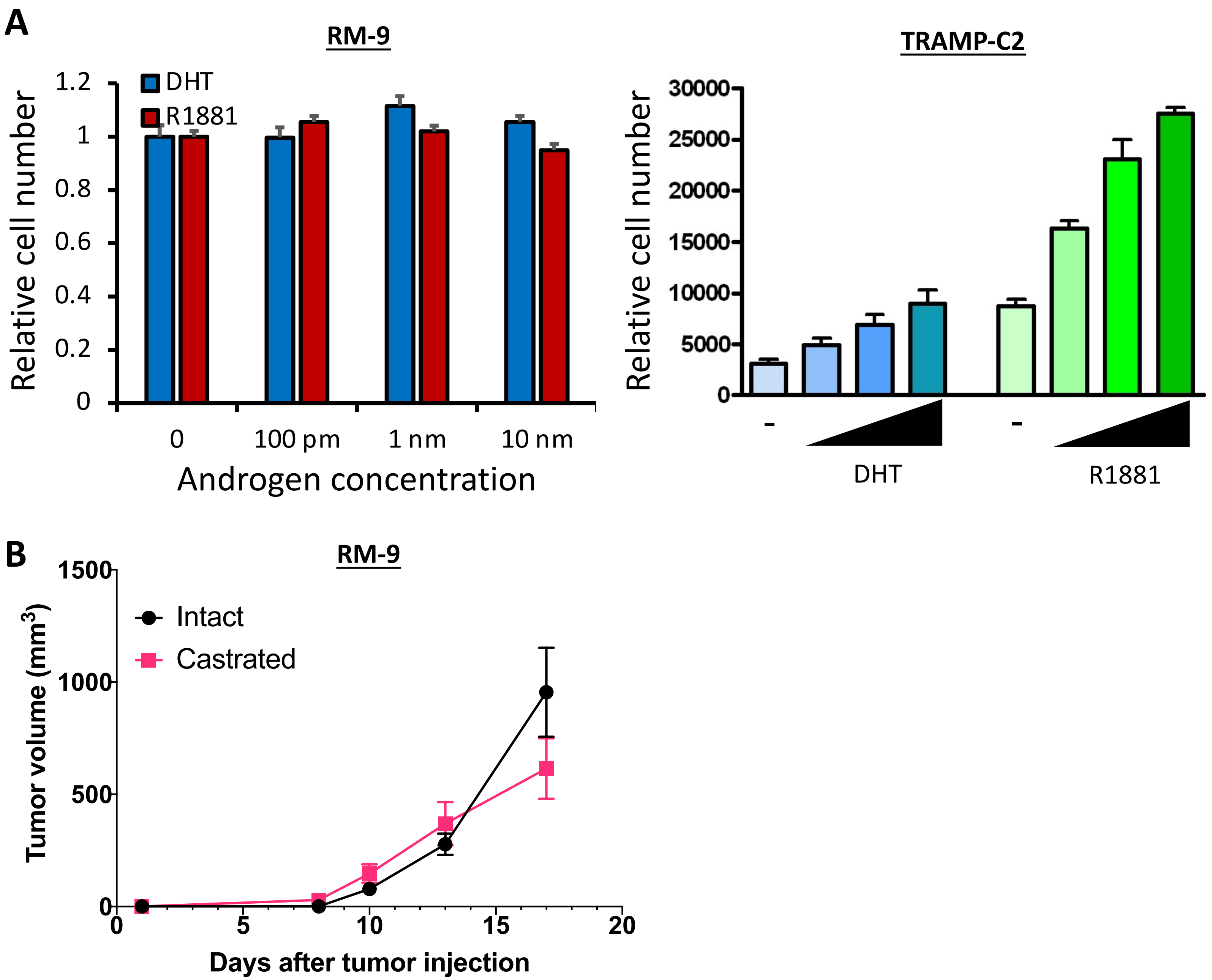


**Supplemental Figure S6: Androgen responsive status of TRAMP-C2 and RM-9 cells.** (A) RM-9 or TRAMP-C2 murine prostate cancer cells were treated with a dose response (0, 100 pM, 1 nM, 10 nM) of androgens (DHT and R1881) for 7 days and relative cell numbers were quantified using a Hochest-based DNA dye. Results are expressed as relative cell number + SE. (B) 50,000 RM-9 cells were subcutaneously injected into intact (black; n=10) or castrated (red; n=7) syngeneic C57BL/6 mice. Tumor growth was monitored by calipers. In our hands, TRAMP-C2 had a low take rate in castrated mice (data not shown).

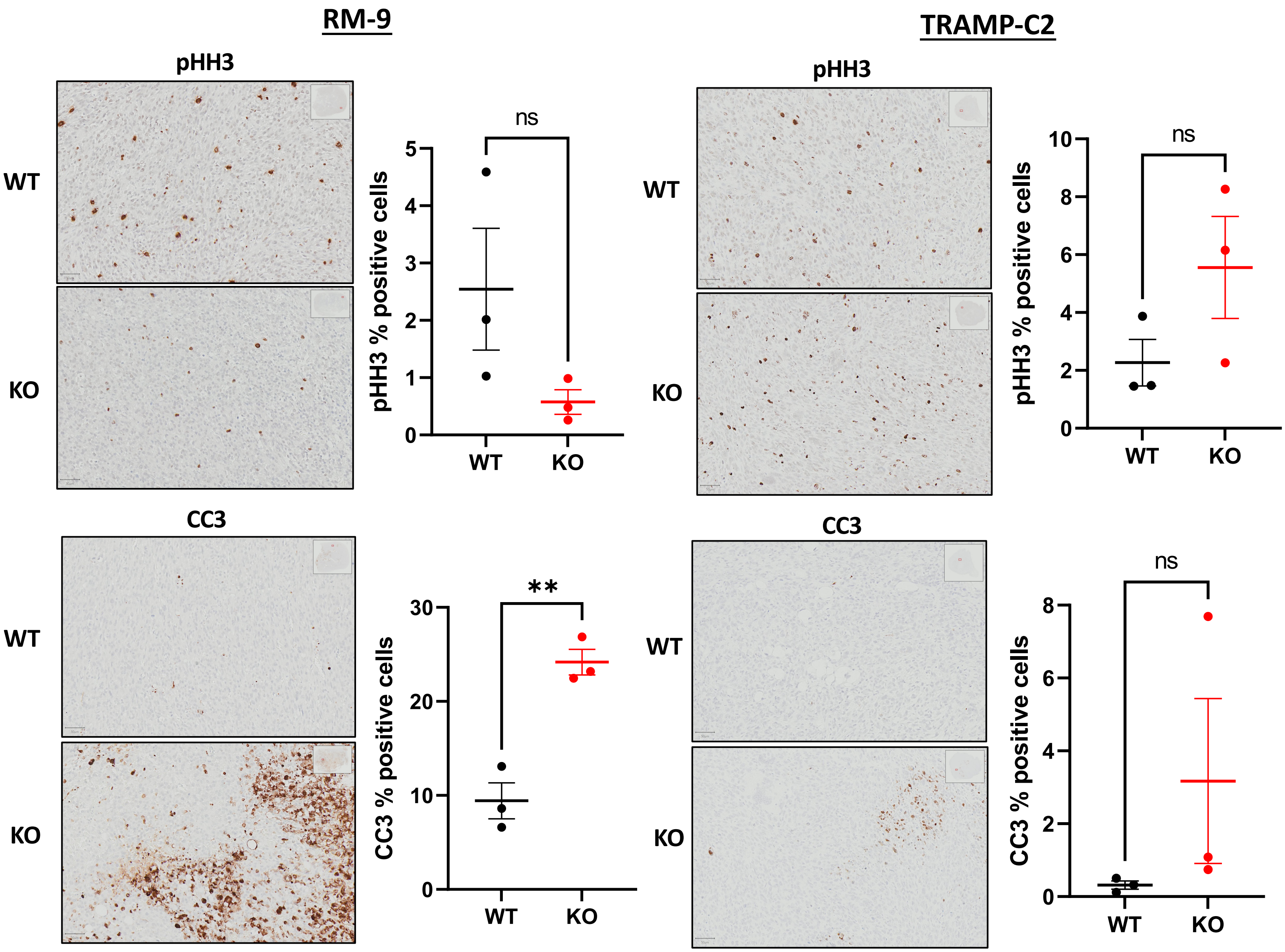


**Supplemental Figure S7: Impact of CAMKK2 status on proliferation and apoptosis in syngeneic mouse models of prostate cancer.** Phospho-histone H3 (Serine-10) (pHH3) and cleaved caspase-3 (CC3) staining in RM-9 (*left*) and TRAMP-C2 (*right*) syngeneic models. ***P* value < 0.01; ns = no significance.
