## Supplemental File 2 for "Systemic ablation of *Camkk2* impairs metastatic colonization and improves insulin sensitivity in TRAMP mice: Evidence for cancer cell-extrinsic CAMKK2 functions in prostate cancer"

Gross anatomy images and weights of livers from TRAMP;*Camkk2*+/+ (WT) and TRAMP;*Camkk2*-/- (KO) mice fed normal diet (ND) or high-fat diet (HFD).

Representative images shown in Figure 3.

### 15-week HFD KO

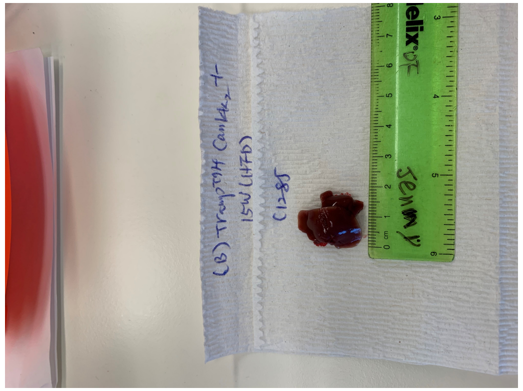
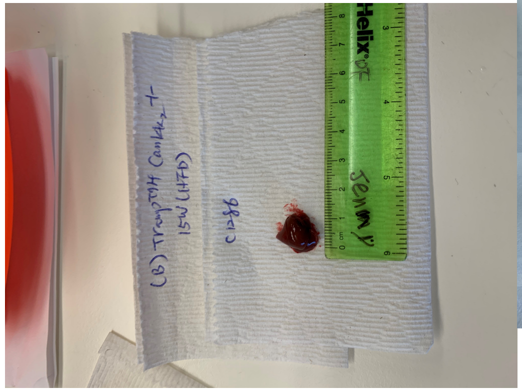
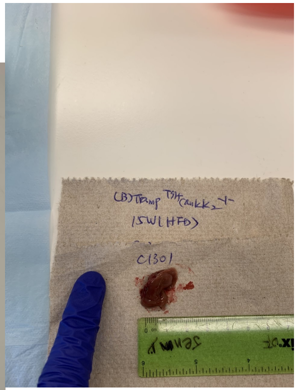
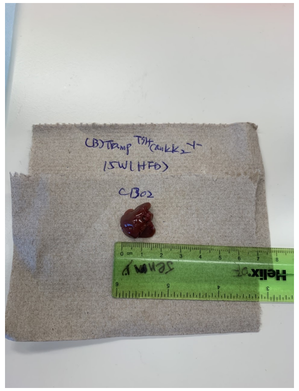
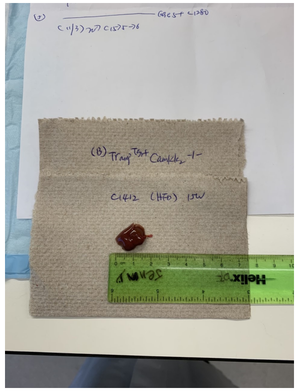
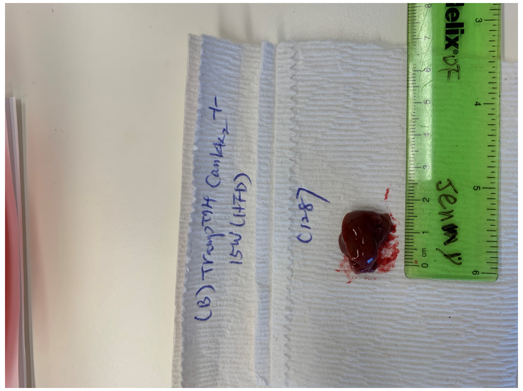
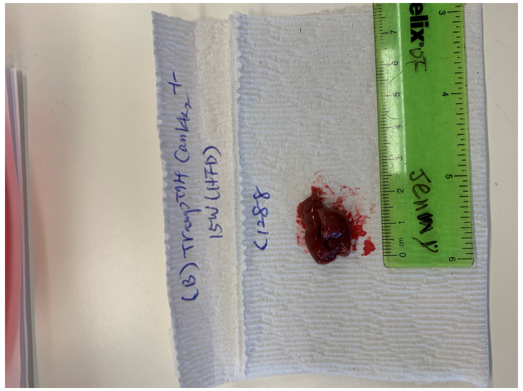
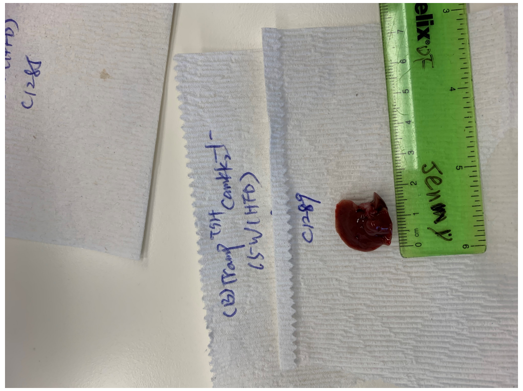

### 15-week HFD WT

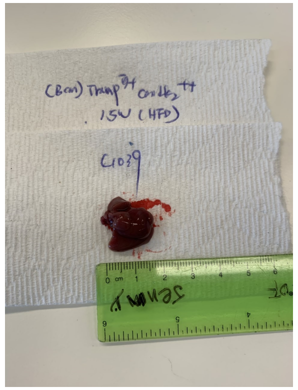
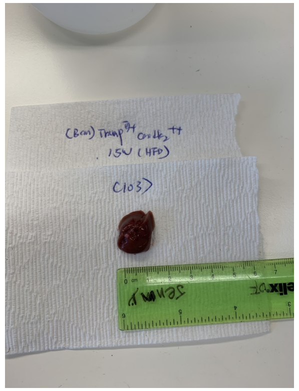
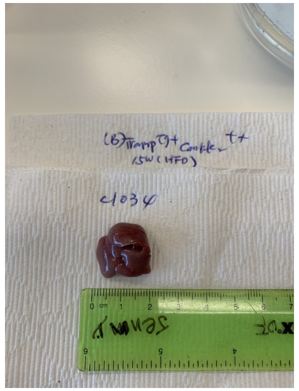
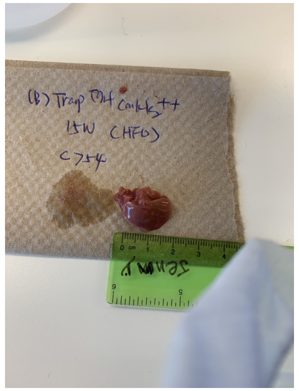
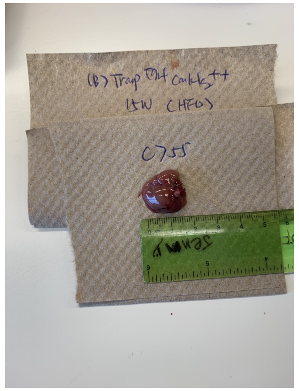
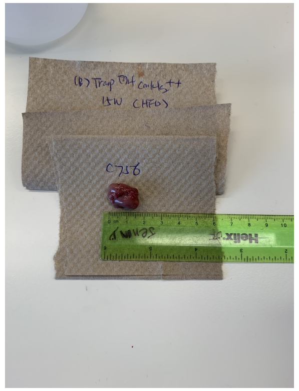
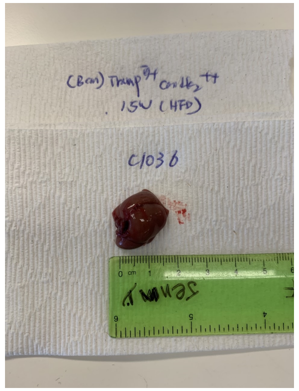
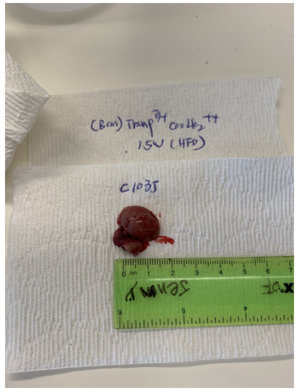
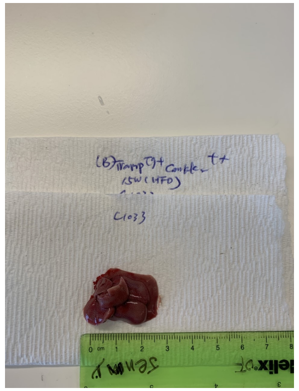

### 30-week HFD KO

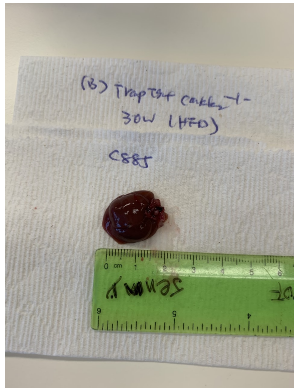

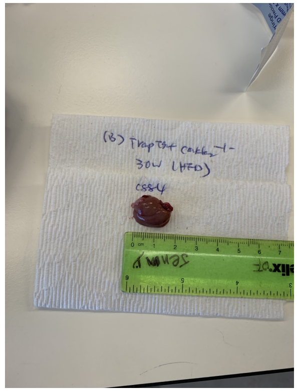

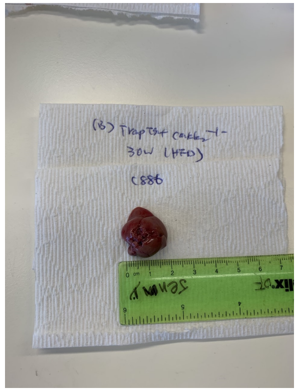

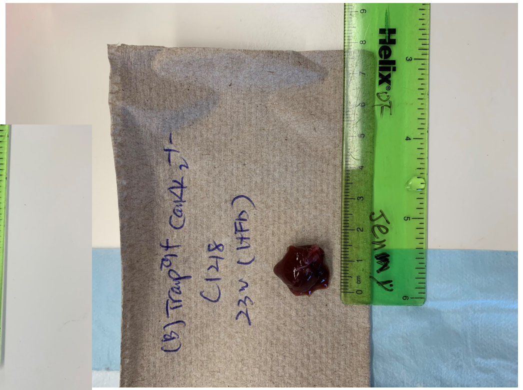

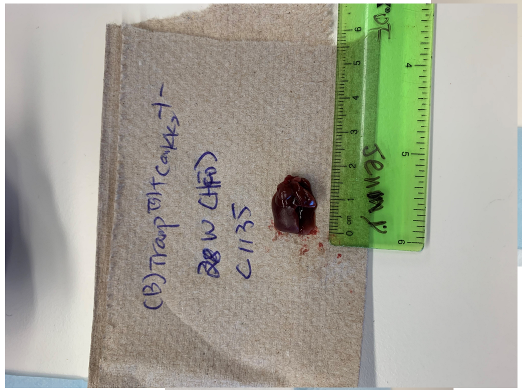

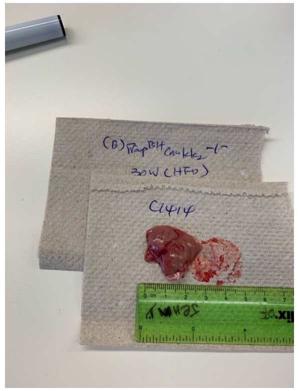

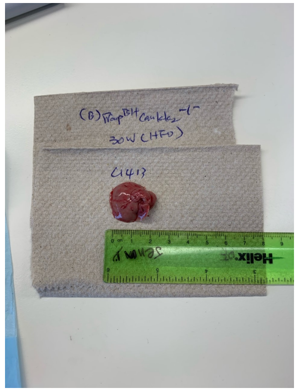

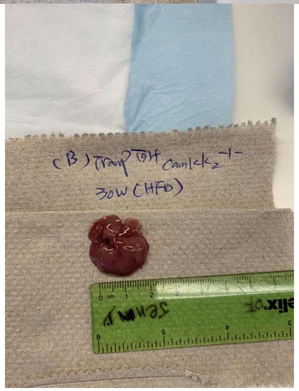

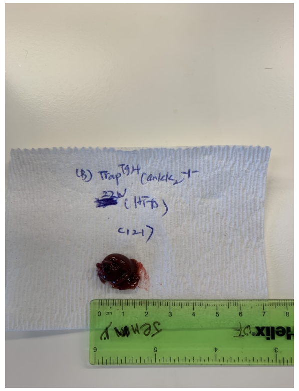

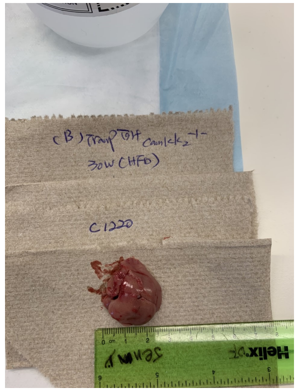

### 30-week HFD WT

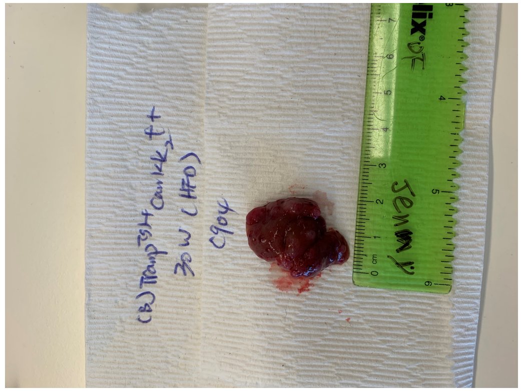


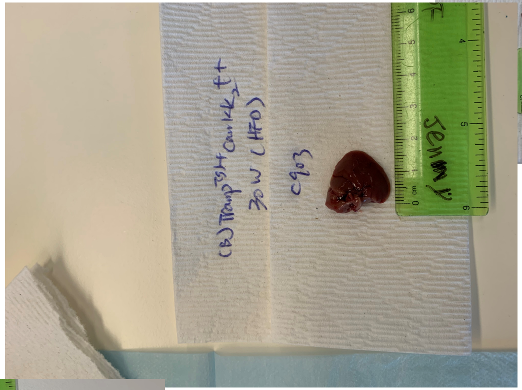


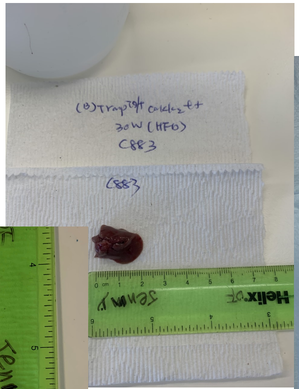


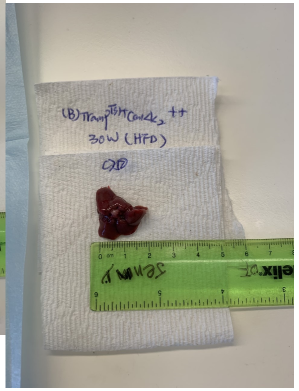

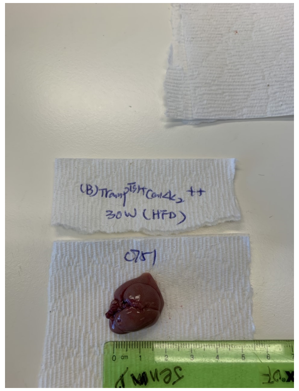


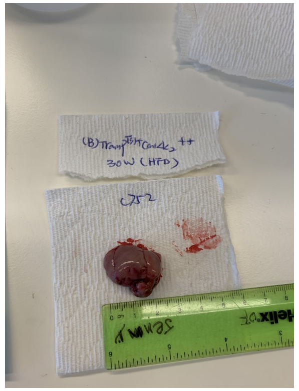


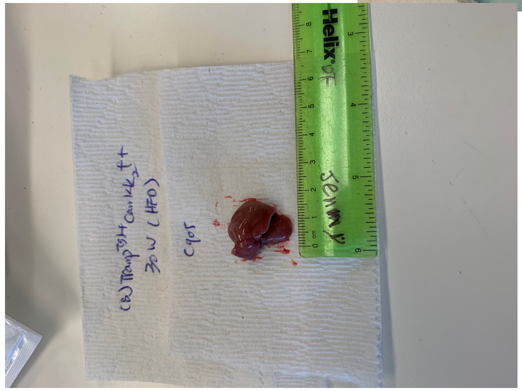


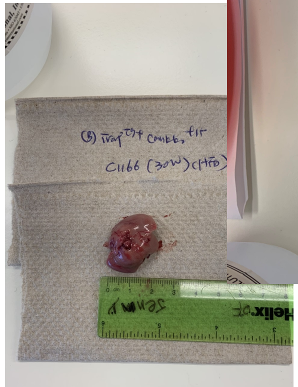


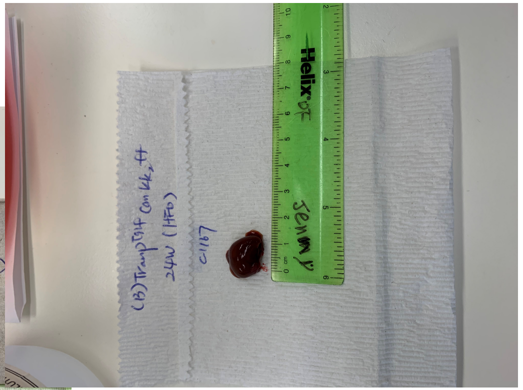


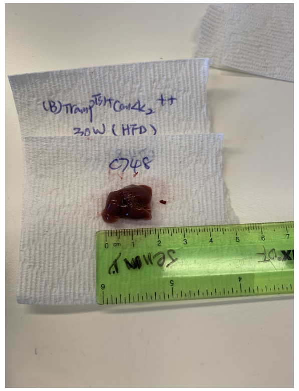


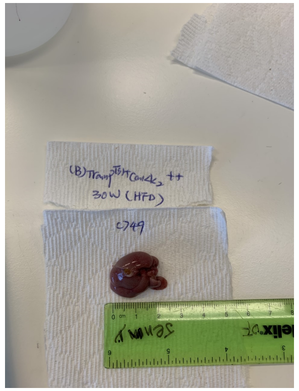

#### Liver weights

| | 15 WKS (HFD) | | | | | | | | | | 30 WKS (HFD) | | | | | | | | | |
|---|---|---|---|---|---|---|---|---|---|---|---|---|---|---|---|---|---|---|---|---|
| **WT** | 1.52 | 1.54 | 1.52 | 2.54 | 1.85 | 1.26 | 1.68 | 1.44 | 1.03 | 1.42 | 1.07 | 2.18 | 1.49 | 2.3 | 3.29 | 2.53 | 1.42 | 2.09 | 3.62 | 1.16 |
| **KO** | 1.51 | 0.7 | 1.38 | 1.43 | 1.22 | 1.27 | 1.2 | 0.99 | | | 1.49 | 2.54 | 1.78 | 0.82 | 1.02 | 1.23 | 2.08 | 3.93 | 2.74 | 2.56 |

| | |
|---|---|
| Average 15wks HFD | WT 1.58 |
| | KO 1.2125 |
